# Chemokines as phosphatidylserine-bound ‘find-me’ signals in apoptotic cell clearance

**DOI:** 10.1101/2020.08.26.269100

**Authors:** Sergio M. Pontejo, Philip M. Murphy

**Author notes:** **Abbreviations** AnV, annexin V; ApoBD, apoptotic bodies; BLI, biolayer interferometry; CL, cardiolipin; DOPC, 1,2-dioleoyl-sn-glycero-3-phosphocholine; DOPE, 1,2-dioleoyl-sn-glycero-3-phosphoethanolamine; DOPEbiot, 1,2-dioleoyl-sn-glycero-3-phosphoethanolamine-N-(cap biotinyl); DOPS, 1,2-dioleoyl-sn-glycero-3-phospho-L-serine; DSPE-PEGbiot, 1,2-distearoyl-sn-glycero-3-phosphoethanolamine-N-[biotinyl(polyethylene glycol)-2000]; ELISA, enzyme-linked immunosorbent assay; FACS, fluorescent-activated cell sorting; GAGs, glycosaminoglycans; GPCR, G protein-coupled receptor; i.p., intraperitoneal; K_D_, binding affinity constant; MFG-E8, milk fat globule epidermal growth factor 8; MVs, microvesicles; oxLDL, oxidized low density lipoproteins; PI, propidium iodide; PS, phosphatidylserine; SN, supernatant; t_1/2_, half life; UV, ultraviolet.

## Abstract

Chemokines are positively charged cytokines that attract leukocytes by binding to anionic glycosaminoglycans (GAGs) on endothelial cells for efficient presentation to leukocyte G protein-coupled receptors (GPCRs). The atypical chemokine CXCL16 has been reported to also bind the anionic phospholipid phosphatidylserine (PS), but the biological relevance of this interaction remains poorly understood. Here we demonstrate that PS binding is in fact a widely shared property of chemokine superfamily members that, like GAG binding, induces chemokine oligomerization. PS is an essential phospholipid of the inner leaflet of the healthy cell plasma membrane but it is exposed in apoptotic cells to act as an ‘eat-me’ signal that promotes engulfment of dying cells by phagocytes. We found that chemokines can bind PS in pure form as well as in the context of liposomes and on the surface of apoptotic cells and extracellular vesicles released by apoptotic cells, which are known to act as ‘find-me’ signals that chemoattract phagocytes during apoptotic cell clearance. Importantly, we show that GAGs are severely depleted from the surface of apoptotic cells and that extracellular vesicles extracted from apoptotic mouse thymus bind endogenous thymic chemokines and activate cognate chemokine receptors. Together these results indicate that chemokines tethered to surface-exposed PS may be responsible for the chemotactic and find-me signal activity previously attributed to extracellular vesicles, and that PS may substitute for GAGs as the anionic scaffold that regulates chemokine oligomerization and presentation to GPCRs on the GAG-deficient membranes of apoptotic cells and extracellular vesicles. Here, we present a new mechanism by which extracellular vesicles, currently recognized as essential agents for intercellular communication in homeostasis and disease, can transport signaling cytokines.

## Introduction

Chemokines comprise a large family of small (8-12 kDa) cytokines that direct leukocyte migration during homeostasis, development, and disease [1]. There are four chemokine subfamilies (C, CC, CXC and CX3C) distinguished and named by the number and arrangement of conserved cysteines near the *N* terminus [2]. In the classic multistep model of leukocyte transendothelial migration, chemokines act by binding sequentially to two main classes of molecules: glycosaminoglycans (GAGs) on endothelial cells and 7-transmembrane domain G protein-coupled receptors (GPCRs) on leukocytes [3]. GAGs act as extracellular scaffolds that concentrate chemokines on the luminal surface of post-capillary venules, positioning them to attract circulating leukocytes by activating specific leukocyte GPCRs [4].

GAGs contain highly anionic sulfated polysaccharides and are thought to protect chemokines from proteases and shear forces in blood vessels [5]. Of note, GAG-binding can induce the formation of high order chemokine oligomers, and chemokine oligomers have a higher GAG binding affinity than chemokine monomers [6]. Thus, GAG-binding and oligomerization may be mutually reinforcing processes that promote formation of haptotactic gradients and presentation of chemokines to leukocytes. Several other types of molecules also bind chemokines; however, the functional significance is not well-defined, and the interactions tend not to be shared in a class sense within or across chemokine subfamilies. A particularly interesting example is CXCL16, which binds the GPCR CXCR6 as well as phosphatidylserine (PS) and oxidized LDL (oxLDL) [7–9]. The latter activities suggested that CXCL16 might function as a scavenger receptor, particularly since it is a multimodular protein containing a transmembrane domain, one of only two chemokines with this type of structure [10]. In fact, CXCL16 was originally named SR-PSOX or ‘<u>s</u>cavenger receptor for <u>PS</u> and <u>ox</u>LDL’ [11]. While the role of the CXCL16-oxLDL interaction in atherosclerosis has been extensively studied [12–14], the biological relevance of PS as a ligand for CXCL16 remains poorly understood. Moreover, although oxLDL has been shown to bind other chemokines [15], to date, CXCL16 is the only reported PS-binding chemokine.

PS is an anionic phospholipid that is normally concealed within the inner leaflet of the plasma membrane of living cells [16]. As cells undergo apoptosis, PS relocates to the outer leaflet of the plasma membrane where it serves as a marker of apoptotic cell death and as an ‘eat-me’ signal promoting recognition and engulfment of apoptotic cells by phagocytes [17–19]. Conversely, chemotactic ‘find-me’ signals, including the chemokine CX3CL1, the nucleotides ATP and UTP, lysophosphatidylcholine, and sphingosine 1-phosphate, guide phagocytes toward apoptotic cells [20–24]. In addition, apoptotic blebs can be shed as PS-containing extracellular vesicles able to recruit phagocytes [25–27]. Cellular receptors for the four classical find-me signals have been identified but how apoptotic blebs can exert their chemotactic action remains unknown. Since most chemokines are highly basic proteins, we hypothesized that anionic phospholipid binding may be a more general property of chemokines. In particular, we considered that chemokines may multitask as apoptotic cell clearance factors tethered to anionic PS on dead cells and apoptotic blebs, just as they are tethered to anionic GAGs on living cells, in both cases to recruit leukocytes via GPCRs.

## Results

### Many human chemokines bind anionic phospholipids including phosphatidylserine

We first conducted a simple screen for chemokine binding to phospholipids using arrays of 15 different phospholipids that are found in biological membranes (left panel in Fig 1A). Of the 10 human chemokines tested, CCL3, CXCL6, CCL5 and CXCL8 displayed very weak or no binding capacity to the array, whereas the other 6 chemokines bound selectively to multiple anionic phospholipids (Fig 1A). The binding pattern was similar for CCL11, CCL21, CXCL9 and CXCL11, with strong binding to PS and cardiolipin (CL), and weaker interaction with the other anionic phospholipids on the array (Fig 1A). CXCL3 also bound to PS and CL, however, its strongest binding was to sulfatide. CCL20 appeared to be CL-selective (Fig 1A). The preference of all 6 of these chemokines for PS and CL over the more highly anionic phosphoinositides suggests some degree of specificity and that these interactions are not solely driven by charge.

**Fig 1.**
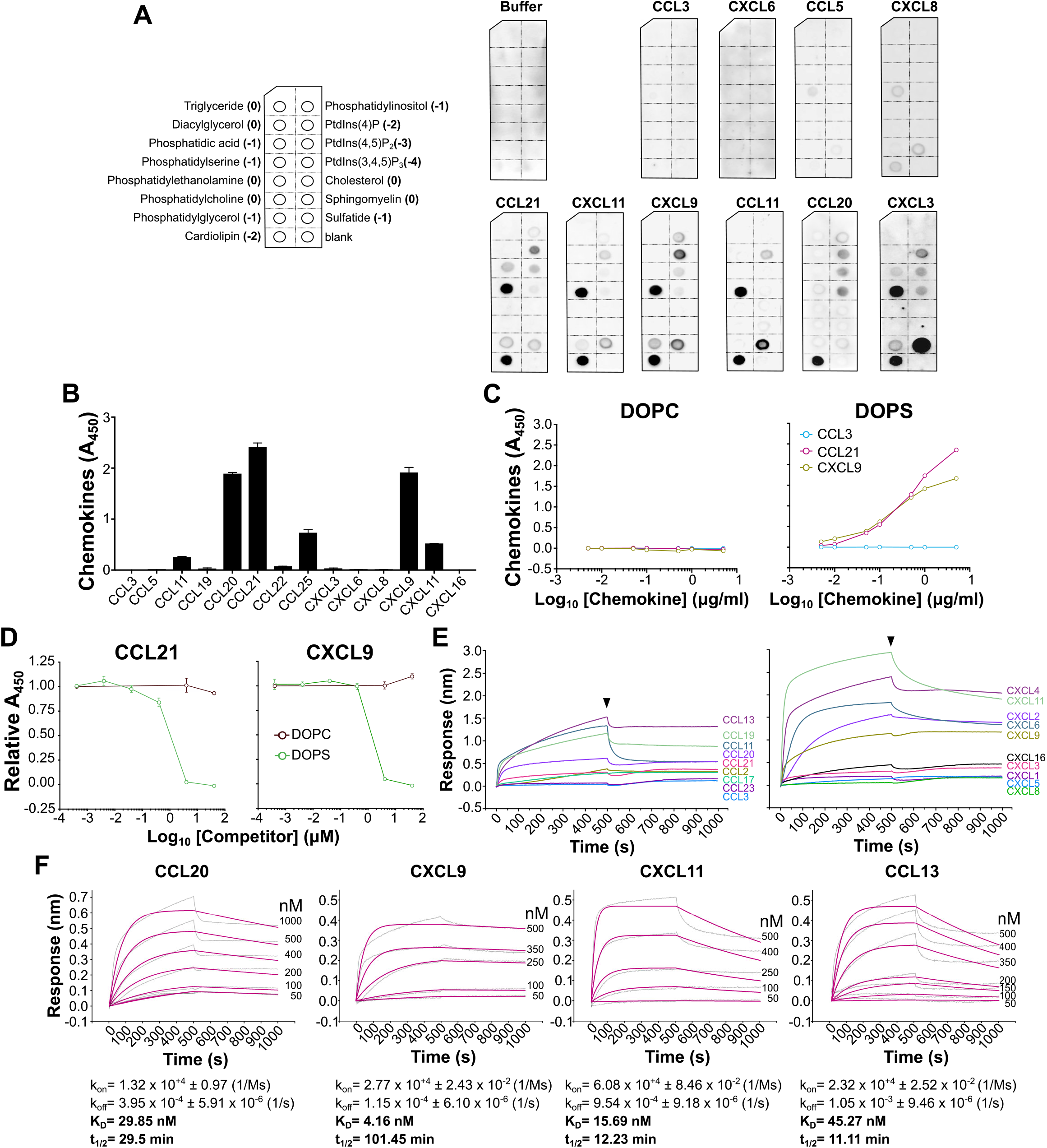
Chemokines bind to anionic phospholipids. The phospholipid-binding activity of chemokines was studied by protein-lipid overlay (A), ELISA (B-D), and BLI (E, F). A) Arrays spotted with 15 different phospholipids (left panel, in parenthesis, net charge of each phospholipid) were incubated with buffer or 0.1 µg/ml of the indicated chemokines. Bound chemokine was detected with specific antibodies. Results are representative of 2-3 experiments for each chemokine. B) Human chemokines indicated on the x-axis were incubated at 1 µg/ml in wells containing immobilized DOPS liposomes. Bound chemokine was detected with specific antibodies for each chemokine. Data are presented as mean ± SD of triplicates in one experiment representative of 2 independent experiments. C) Increasing concentrations of CCL3, CCL21 or CXCL9 were incubated in wells containing immobilized DOPC (left panel) or DOPS (right panel) liposomes. Bound chemokine was detected as in B. Data are presented as mean ± SD of triplicates from one experiment representative of 3 independent experiments. D) Increasing concentrations of the soluble phospholipids indicated in the inset of the right panel were preincubated with 0.5 µg/ml of CCL21 or CXCL9. The chemokine-lipid mix was then added into wells containing immobilized DOPS liposomes, and liposome-bound chemokine was detected by specific antibodies. Data are presented as mean ± SD of triplicates in one experiment representative of 3 independent experiments. E) Chemokine-PS binding screen by BLI. Biosensors immobilized with DOPS liposomes were incubated with 1 µM of the indicated human CC (left panel) and CXC (right panel) chemokines color-coded on the right side of each curve. The binding response in nm (y-axis) over time (x-axis) for each chemokine is shown. The binding of each chemokine to biosensors coated with DOPC liposomes was used as reference and subtracted from the corresponding binding curve. Black arrowheads point to the beginning of the dissociation phase. F) Kinetic analysis. Selected chemokines were incubated at the indicated concentrations with biosensors coated with DOPS liposomes. Binding of each chemokine concentration to DOPC biosensors was subtracted. Curves were globally fitted to a 1:1 Langmuir model and the association (k_on_), dissociation (k_off_) and affinity (K_D_) constants were calculated. The half-life (t_1/2_) for each interaction was calculated as t_1/2_=ln(2)/k_off_.

Since CL is not found in the plasma membrane of mammalian cells, we focused on the chemokine-PS interaction. To confirm that chemokines can interact with PS in the context of a phospholipid bilayer, we studied by ELISA chemokine binding to PS-containing liposomes. For this, we immobilized DOPC liposomes (containing phosphatidylcholine [PC] only) or DOPS liposomes (containing 30% PS) containing trace amounts of biotinylated DOPE (DOPEbiot) onto streptavidin-coated plates. See ‘Materials and Methods’ for the exact composition of these liposomes and definition of phospholipid acronyms. Consistent with the results in Fig 1A, CCL11, CXCL11, CCL21 and CXCL9, but not CCL3, CCL5, CXCL6 and CXCL8 bound to DOPS liposomes to some extent (Fig 1B). Of note, since different primary antibodies were used to detect each chemokine, direct quantitative comparisons cannot be made from this ELISA-based lipid-binding experiment. Importantly, CXCL9 and CCL21, but not CCL3, bound to DOPS but not to DOPC liposomes in a dose-dependent manner (Fig 1C), and increasing concentrations of soluble DOPS (IC_50_ ≈ 1 µM) but not DOPC blocked their binding to DOPS liposome-coated plates (Fig 1D). These results confirmed the PS-specificity of these chemokines. On the other hand, as shown in Fig 1B, CXCL3 was not able to recognize PS in liposomes, and CCL20, which appeared to be CL-specific by lipid array (Fig 1A), displayed a strong binding to DOPS liposomes by ELISA. Besides the different disposition of the lipids (spotted purified lipids vs liposomes), differences in the experimental conditions may account for these few discrepancies between lipid array- and ELISA-based binding experiments. Also, unexpectedly, we did not detect binding of CXCL16 to DOPS liposomes by ELISA (Fig 1B).

To directly and reliably compare the PS-binding activity of different chemokines without the need of exogenous protein tags or detection antibodies, we next analyzed chemokine-PS binding by biolayer interferometry (BLI). To validate BLI as a method to study liposome-protein interactions, we first analyzed binding of milk fat globule-epidermal growth factor 8 (MFG-E8 or lactadherin), a well-known PS-binding protein [28]. As shown in S1 Fig, MFG-E8 bound to biosensors immobilized with DOPS liposomes, but not with DOPC liposomes. Using this technique, we analyzed PS-binding for 21 human chemokines (S1 Table). As shown in Fig 1E and S1 Table, CXCL16 but also many other chemokines from the CC and CXC subfamilies, including CXCL3, whose binding was not detectable by ELISA (Fig 1B), bound to PS-containing liposomes. Consistent with the ELISA and phospholipid array results, CCL3 and CXCL8 did not interact with DOPS liposomes by BLI, but CXCL6, which lacked PS binding activity in previous experiments, strongly bound DOPS liposomes by BLI (Fig 1E). Besides the numerous differences in the experimental conditions, the few discrepancies observed between BLI and the antibody-based binding assays might be due to the inability of certain anti-chemokine antibodies to detect their target chemokine complexed with lipid. To test this hypothesis we analyzed by BLI the binding of CCL21 and CXCL9, which consistently bound to PS in all experimental settings, and of CXCL6 and CXCL3, which only bound DOPS liposomes by BLI, to their corresponding detection antibodies in the presence of DOPS liposomes. In BLI experiments, at the same concentration, large analytes (chemokine-liposome complex) are expected to cause larger binding responses than small molecules (chemokine alone). As shown in S2 Fig, binding of CCL21 and CXCL9 to BLI biosensors immobilized with the appropriate anti-chemokine antibody was increased in the presence of DOPS liposomes but not DOPC liposomes. In contrast, this signal increase was not observed for CXCL6 and CXCL3 (S2 Fig), suggesting that the corresponding antibodies are unable to detect these chemokines in a complex with liposomes. This may explain why the PS-binding of CXCL6 and CXCL3 was not detectable by ELISA. Also, importantly, the interaction of most chemokines with PS appeared to be very stable, as indicated by a slow dissociation phase (Fig 1E). We determined the kinetic parameters for 4 selected chemokines and obtained binding affinity constants (K_D_) ranging from 4-45 nM and half-life (t_1/2_) values between 10 and 100 min (Fig 1F). This binding affinity is near the range of the affinity of the MFG-E8-PS interaction (K_D_ = 3.3 nM) [29]. These binding and kinetic analyses were performed with liposomes containing 30% PS but we found that the PS detection limit of CCL20, CCL19 and CXCL11 was in the 5-10% range (S3 Fig). Taken together, these results demonstrate that CXCL16 is not exceptional since PS is a ligand for many human chemokines.

### Anionic phospholipids induce chemokine oligomerization

Chemokines bound to cell surface GAGs oligomerize and form haptotactic gradients. We tested whether PS or other phospholipids were also capable of inducing chemokine oligomerization. Using cross-linking assays we found that, consistent with their lipid-binding specificity, CCL11, CXCL11, CXCL9 and CCL21 oligomerized in the presence of either PS or CL, but not in the presence of phosphatidylethanolamine (PE) or PC (Fig 2). In contrast, the oligomeric state of CCL3, which does not interact with any of these phospholipids, was unaltered by PS and CL (Fig 2). These results further confirmed the interaction of chemokines with anionic phospholipids and support PS as a new potential substrate for the formation of chemokine oligomers and haptotactic gradients.

**Fig 2.**
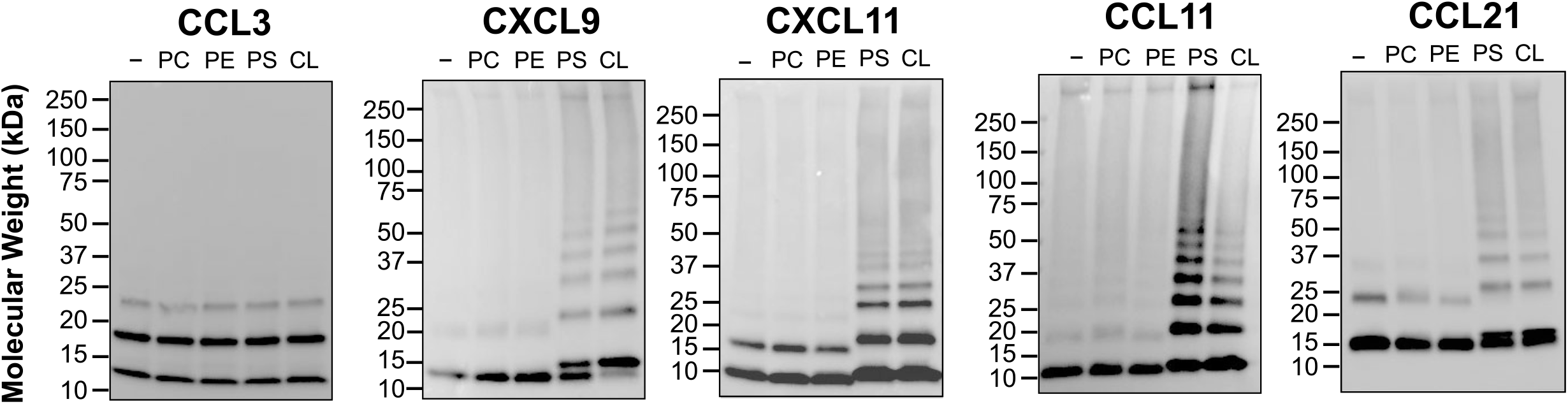
Anionic phospholipids induce oligomerization of phosphatidylserine-binding chemokines. The chemokines (50 ng) indicated above each panel were incubated in the absence or presence of the phospholipids (1:8, chemokine:lipid molar ratio) indicated above each lane (-, no lipid; PC, phosphatidylcholine; PE, phosphatidylethanolamine; PS, phosphatidylserine; CL, cardiolipin) with the crosslinker BS_3_. Samples were analyzed by SDS-PAGE and immunoblot using specific anti-chemokine antibodies. Molecular weight markers are shown in kDa on the left of each panel. Data are representative of 2 independent experiments for each chemokine.

### Chemokine binding to phosphatidylserine does not interfere with chemotactic activity

Next, we investigated whether chemokine binding to DOPS liposomes affects chemokine bioactivity. For this, we evaluated two PS-binding chemokines, CCL20 and CCL21, and included the non-binder CCL3 as a negative control. Each chemokine was tested for its ability to chemoattract L1.2 mouse B cell lymphoma cells expressing the cognate mouse receptors Ccr6, Ccr7 and Ccr1, respectively, in the presence of increasing doses of DOPS or DOPC liposomes. As shown in the left column of Fig 3A, 1 nM CCL20, CCL21 and CCL3 induced potent chemotactic responses that remained unaltered even in the presence of a 10^5^-fold molar excess of DOPS liposomes. As expected, DOPC liposomes did not inhibit any of these chemokines (Fig 3A). In order to confirm that the chemokines bound to the liposomes under these experimental conditions, the presence of liposome-bound chemokine was analyzed by ELISA. Using the DOPEbiot introduced in our liposomes, a small aliquot of each chemokine-liposome mix prepared for the chemotaxis assays was immobilized on streptavidin-coated plates. Consistent with their PS-binding activity, CCL20 and CCL21 but not CCL3 were detected only when mixed with DOPS-liposomes (right column in Fig 3A). These findings indicate that high doses of PS do not impair the activity of PS-binding chemokines, but they do not directly prove whether or not a chemokine-liposome complex can activate the receptor. To address this question, we analyzed the chemotactic activity of CCL20 alone or mixed with DOPS or DOPC liposomes after pull-down with Strep-Tactin (streptavidin analog)-coated beads. As shown in panel A of S4 Fig, pull-down with Strep-Tactin-beads markedly reduced the levels of CCL20-DOPS complexes in solution, which resulted in at least a 40% reduction of cell migration compared to pull-down supernatants from samples containing CCL20 alone or CCL20 + DOPC liposomes (panel B in S4 Fig). In contrast, pull-down did not affect migration induced by the non-PS-binding chemokine CCL3 in the presence of DOPS liposomes (panel B in S4 Fig). These results suggest that the CCL20-DOPS complex contributes to the overall cell migration observed in Fig 3A, and therefore, at least in the case of CCL20, PS-binding appears to be compatible with the chemotactic activity of the chemokine. This observation contrasts with the inhibitory effect reported for soluble GAGs on several chemokines and the favored notion that while GAGs present chemokines on endothelial cells, they may need to be released to act at leukocyte GPCRs [30,31]. In fact, we found that while a 10^4^-fold molar excess of DOPS or DOPC liposomes did not alter the chemotactic potency or the efficiency of CCL3, CCL20 and CCL21, a 10^3^-fold molar excess of soluble heparin sufficed to nearly neutralize CCL20 and CCL21 (Fig 3B). Conversely, heparin did not block CCL3 (Fig 3B), probably due to the known low binding affinity of this chemokine for GAGs [6,32]. Thus, PS-binding chemokines engage PS by a molecular mechanism that, unlike GAG engagement, permits binding of the chemokine to cellular receptors.

**Fig 3.**
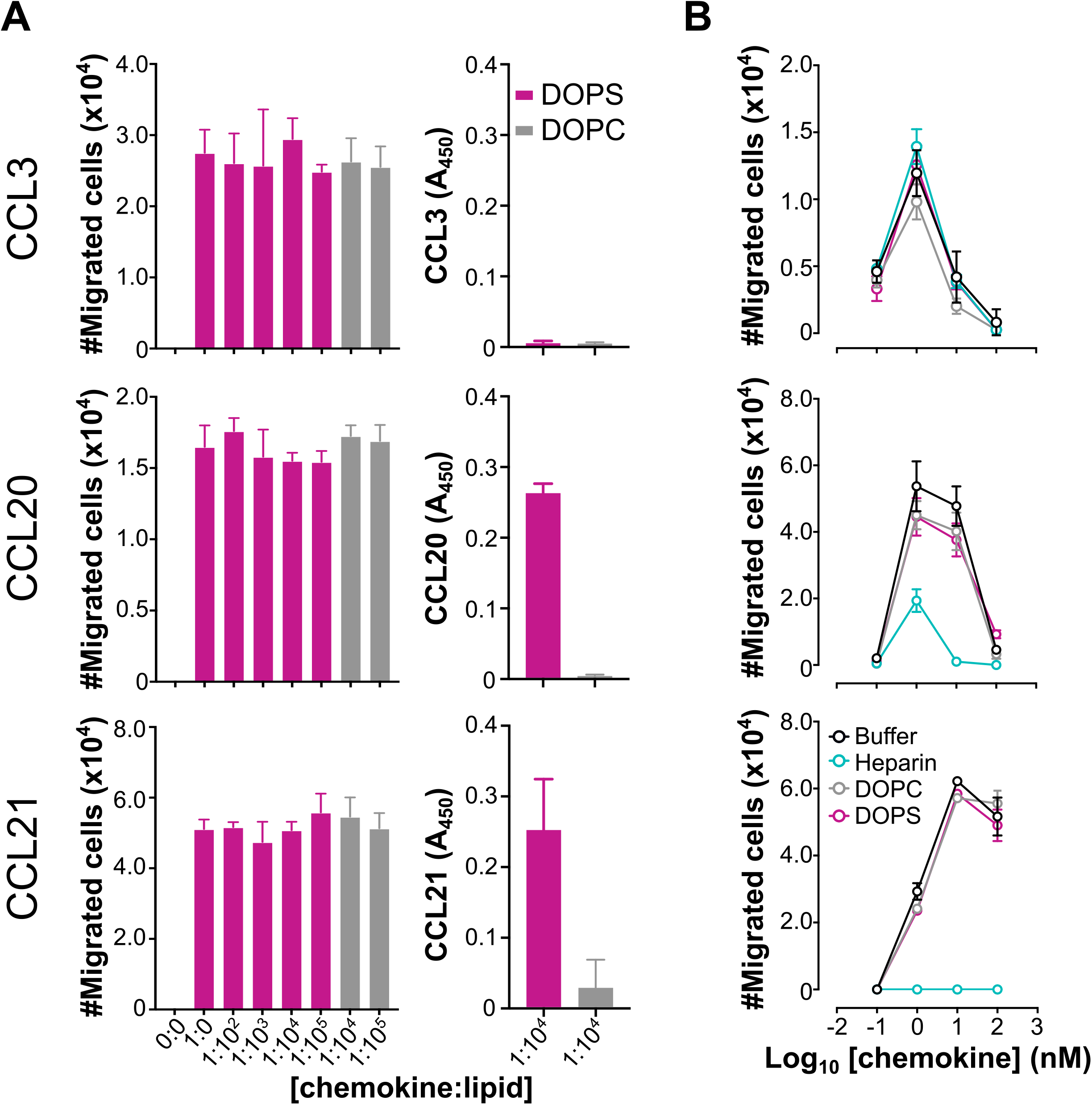
Chemokine interaction with liposomal phosphatidylserine does not affect chemotactic activity. A) *Left column*, DOPS liposomes do not impair chemokine chemotactic activity. The indicated chemokines (1 nM) were preincubated for 30 min at room temperature with the indicated molar excess (x axis) of DOPS (magenta) or DOPC (gray) liposomes. Then, chemotactic activity of the mixture was tested using L1.2 reporter cells stably expressing the mouse receptors Ccr1 (for CCL3), Ccr6 (for CCL20) and Ccr7 (for CCL21). Bars correspond to the total number of migrated cells after 3-4 h at 37°C for each chemokine:lipid mix or buffer alone (0:0). *Right column*, CCL20 and CCL21 but not CCL3, bind to DOPS liposomes in the experimental conditions used for the chemotaxis assays. Detection of liposome-bound chemokine by ELISA. 50 µl of the 1:10^4^ molar ratio mix of each chemokine (as indicated on the left side of each row) with DOPS liposomes or DOPC liposomes (as indicated in the inset of the top graph) were incubated in streptavidin-coated wells. After washing, liposome-bound chemokine was detected by specific antibodies. All data are presented as mean ± SD of triplicates from one experiment representative of 3 independent experiments. B) Anionic GAGs but not PS-containing liposomes inhibit the chemotactic activity of CCL20 and CCL21. Increasing concentrations (0.1 to 100 nM) of CCL3, CCL20 and CCL21 (as indicated on the left side of each row) were preincubated for 30 min at room temperature with buffer alone (black), or a 10^4^-fold molar excess of DOPC liposomes (gray) or DOPS liposomes (magenta), or a 10^3^-fold molar excess of heparin (blue), as indicated in the inset of the bottom graph. Chemotactic activity for each condition is presented as the mean ± SD number of migrated cells of triplicates from one experiment representative of 2-3 independent experiments.

### Phosphatidylserine-binding chemokines interact with the surface of apoptotic cells and apoptotic blebs

Since PS is normally located on the inner leaflet of the plasma membrane of living cells but flips to the outer leaflet of the plasma membrane of cells undergoing apoptosis [16], apoptotic cell death is an obvious biological process in which PS-binding by chemokines may occur and be functionally relevant. This property of PS is exploited for detection of apoptotic cells by staining with the PS-binding protein annexin V (AnV) [33]. Similarly, apoptotic blebs released by dying cells also expose PS on their surface [34]. Therefore, we studied whether chemokines can interact with the surface of apoptotic blebs, apoptotic CHO cells and apoptotic mouse primary thymocytes.

In order to obviate background interference by cell surface GAGs, we initially used GAG-deficient CHO-745 cells rendered apoptotic by ultraviolet (UV) light irradiation. After treatment, two distinct cell populations were observed by fluorescence-activated cell sorting (FACS), FSC high (FSC^hi^) and low (FSC^lo^), with the percentage of FSC^lo^ cells being markedly increased after exposure to UV light (Mock FSC^lo^, 10.3%; UV FSC^lo^, 34.7%) (Fig 4A). In addition, a high percentage (19.8%) of small SSC low (SSC^lo^) events, most likely corresponding to apoptotic blebs, was detected in UV-treated CHO-745 cells (Fig 4A). We found that most FSC^lo^ events were AnV^+^ and propidium iodide^+^ (PI^+^) (late apoptotic), whereas FSC^hi^ cells were AnV^−^ PI^−^ (non-apoptotic, or live) (Fig 4A). Therefore, for subsequent chemokine binding experiments, we relied on this FSC profile to distinguish live and late apoptotic cells without the need of AnV staining, which might interfere with chemokine binding to cell surface-exposed PS. We analyzed the binding of 4 biotinylated proteins, AnV as positive control, CCL3 as negative control and two PS-binding chemokines, CCL11 and CXCL11, to 3 sources of target membranes: FSC^lo^ late apoptotic cells, FSC^hi^ live cells and SSC^lo^ apoptotic blebs. As shown in Fig 4B, the PS-binding proteins, AnV, CCL11 and CXCL11, but not CCL3, displayed a ≥2-log stronger binding to FSC^lo^ cells than to FSC^hi^ cells from either mock- or UV-treated cells. Of note, like AnV, CCL11 and CXCL11 similarly bound Mock-FSC^lo^ and UV-FSC^lo^ cells, indicating that dying cells spontaneously generated in the mock sample and those induced by UV irradiation were equivalent regarding their PS exposure in the plasma membrane. Furthermore, we observed that CCL11, CXCL11 and AnV, but not CCL3, bound to the surface of PI^lo^ apoptotic blebs (Fig 4C). Therefore, we concluded that PS-binding chemokines interact with the surface of late apoptotic cells and apoptotic blebs. In addition to these commercial synthetic non-glycosylated chemokines, we confirmed that glycosylated CCL21 and CXCL9 produced in Expi293F cells also bound to PS-containing liposomes and apoptotic CHO-745 cells (S5 Fig). To rule out the possibility that the cognate cellular receptors of CCL11, CXCL9, CXCL11 and CCL21 could be mediating their binding to apoptotic cells, we analyzed the expression of CCR3 (receptor for CCL11), CXCR3 (receptor for CXCL9 and CXCL11) and CCR7 (receptor for CCL21) in CHO-745 cells. As shown in panel A of S6 Fig, no significant expression of these receptors was detected in either mock-treated or AnV^−^ and AnV^+^ populations in UV-irradiated cells. Thus, PS-binding chemokines interact with apoptotic cells and blebs in a GAG- and chemokine receptor-independent manner.

**Fig 4.**
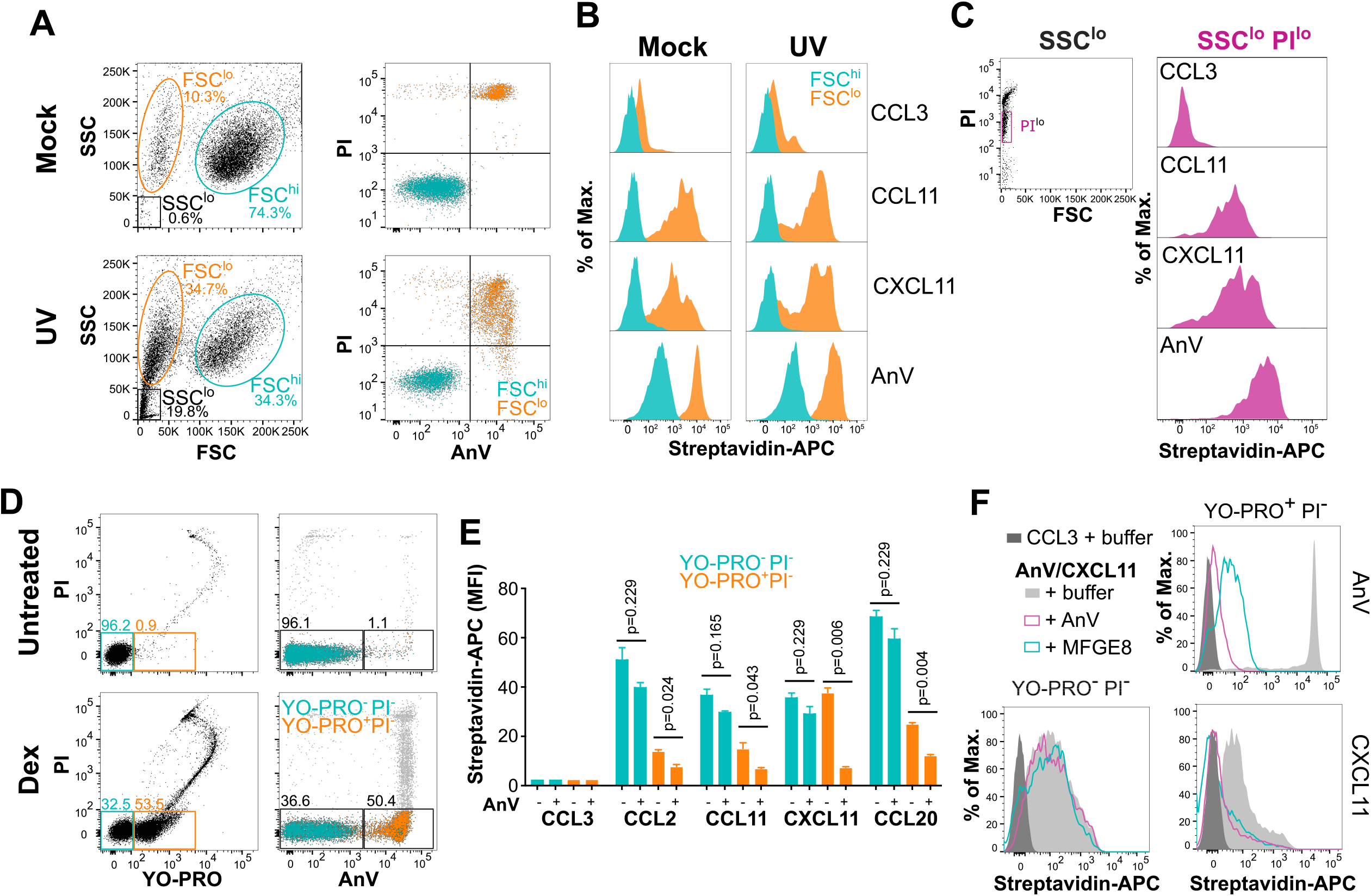
Phosphatidylserine-binding chemokines bind to the surface of apoptotic cells and apoptotic blebs. (A-C) GAG-deficient CHO-745 cells. (D-F) Mouse thymocytes. A) FACS analysis and gating of mock- and UV-irradiated CHO-745 cells. In the left column, FSC-SSC dot plots. Gates for FSC^lo^, FSC^hi^ and SSC^lo^ populations are shown in orange, blue, and black, respectively. In the right column, dot plots for annexin V (AnV) and propidium iodide (PI) staining of FSC^lo^ and FSC^hi^ populations appropriately colored. B and C) binding histograms of the indicated biotinylated proteins to FSC^lo^ (blue) and FSC^hi^ (orange) cells from mock and UV-irradiated cells in B, or to SSC^lo^ events (magenta) in C. In C, *left graph*, PI-FSC dot plot showing the gate of PI^lo^ events included in the binding analysis in the right column. Protein binding was detected with streptavidin-APC (x axis). D) PI-YO-PRO (left column) and PI-AnV (right column) dot plots of freshly isolated mouse thymocytes (untreated) or incubated ex vivo with 1 µM dexamethasone (Dex). In the left column, gates for YO-PRO^−^PI^−^ (blue, live cells) and YO-PRO^+^PI^−^ (orange, apoptotic cells) cells are shown, and their AnV-PI profile is represented in the right column with events color-coded. Numbers above each gate indicate % of total events. E) Quantification of the median fluorescence intensity (MFI) of the binding of the indicated biotinylated chemokines to live cells (YO-PRO^−^PI^−^, blue) or early apoptotic cells (YO-PRO^+^PI^−^, orange) from Dex-treated mouse thymocytes in the absence or presence of unlabeled AnV as indicated on the x-axis. Bars represent mean ± SD of duplicates from one experiment representative of 3 independent experiments. p values from multiple t tests with Holm-Sidak correction for multiple comparisons are indicated. F) Binding histograms of CCL3 (solid dark gray), or CXCL11 and AnV (as indicated on the right side of each graph row) in the presence of buffer (solid light gray) or unlabeled AnV (open magenta) and MFG-E8 (open blue), as indicated in the legend, to live (YO-PRO^−^PI^−^) or early apoptotic cells (YO-PRO^+^PI^−^) as indicated above each graph column. In E and F, binding of biotinylated proteins was detected with streptavidin-APC.

UV irradiation of CHO-745 cells failed to produce AnV^+^ PI^−^ cells, a population typically termed “early apoptotic cells” [35]. To study the binding of chemokines to early apoptotic cells while at the same time expanding our analysis to primary cells, we induced apoptosis in mouse thymocytes by ex vivo treatment with dexamethasone, a well characterized model of apoptosis in primary lymphocytes [36]. To distinguish live from apoptotic cells without using AnV, which might interfere with chemokine binding to PS, we used YO-PRO, which selectively permeates apoptotic cells in a PS-independent manner [37]. As shown in Fig 4D, 4h after dexamethasone treatment, YO-PRO and AnV detected similar levels of early apoptotic thymocytes (53.5 and 50.4%, respectively). Then, we studied the binding of CCL3 (negative control) and 4 PS-binding chemokines, CCL2, CCL11, CXCL11 and CCL20, to live (YO-PRO^−^ PI^−^) and early apoptotic (YO-PRO^+^ PI^−^) thymocytes. CCL2, CCL11, CXCL11 and CCL20, but not CCL3, bound to both live and apoptotic cells (Fig 4E). Importantly, in the presence of unlabeled AnV, the binding of these chemokines to apoptotic cells was reduced by at least 50% whereas AnV inhibition of chemokine binding to live cells was trivial (Fig 4E). In particular, CXCL11 binding to apoptotic cells was nearly completely abrogated not only by AnV but also by MFG-E8 (Fig 4E and 4F). Also, as expected, unlabeled AnV and MFG-E8 highly reduced the binding of biotinylated AnV to apoptotic cells (Fig 4F). In contrast, these two PS-binding proteins did not affect the interaction of CXCL11 with live cells (Fig 4E and 4F). This demonstrates that CXCL11 binding to apoptotic cells but not to live cells is PS-mediated. Unlike CHO-745 cells, thymocytes may contain surface GAGs that may contribute to the chemokine binding to the cell surface. By analyzing cell binding activity for the strong GAG-binding protein B18 from poxvirus [38], we confirmed the presence of GAGs in live (AnV^−^ PI^−^) thymocytes (Panel A in S7 Fig). However, interestingly, B18 binding was massively downregulated on apoptotic thymocytes (AnV^+^ PI^−^) (Panel A in S7 Fig). Consistent with this, we found that B18 binding to UV-irradiated GAG-competent CHO-K1 cells was markedly reduced even when these cells were still negative for AnV staining (Panel B in S7 Fig). These observations indicate that while cell surface GAGs could mediate chemokine binding to live cells, they are not involved in the binding to at least these two types of apoptotic cells. Like GAGs, cellular chemokine receptors might also mediate chemokine binding to the surface of thymocytes. We found low expression levels of Ccr2 and Ccr6 (receptors for CCL2 and CCL20, respectively) but high expression of Ccr3 (receptor for CCL11) on apoptotic (AnV^+^) thymocytes but not on live (AnV^−^) thymocytes (Panel B in S6 Fig). In contrast, apoptotic thymocytes did not express Cxcr3, the receptor for CXCL11 (Panel B in S6 Fig). This may explain why AnV neutralized CXCL11 binding to apoptotic cells, yet only partially blocked CCL2, CCL20 and CCL11 binding. In summary, we conclude that PS-binding chemokines can bind to early apoptotic cells by a mechanism involving surface-exposed PS.

### Active endogenous phosphatidylserine-binding chemokines are transported on the surface of extracellular vesicles from apoptotic mouse thymus

We have demonstrated that many exogenous recombinant human chemokines are able to bind PS presented in pure form or on liposomes or on the surface of apoptotic blebs and apoptotic cell lines and primary cells. Next, we analyzed whether this interaction occurs in vivo with endogenous chemokines by inducing apoptosis and the production of apoptotic blebs in the thymus of C57BL/6j mice by i.p. injection of dexamethasone.

Although we did not detect a high number or frequency of apoptotic thymocytes in dexamethasone-treated mice, possibly due to rapid elimination by phagocytes, the high number of apoptotic blebs observed in these mice provided evidence for the efficacy of the treatment (Panel A in S8 Fig). As expected, the apoptotic blebs exposed PS on their surface as indicated by AnV^+^ staining (Panel B in S8 Fig). Using antibody-based arrays, we identified 6 chemokines that were highly expressed or induced in thymus extracts 6 h and 18 h after dexamethasone treatment: Ccl21, Ccl6, Ccl12, Cxcl10, Ccl9/10 and Ccl5 (Fig 5A). Of these, only Ccl12, Ccl21 and Cxcl10 bound to DOPS liposomes (Fig 5B). Therefore, these 3 chemokines were the main candidates to engage with PS on the surface of extracellular vesicles from mouse thymus. To test this, we first followed a previously reported protocol to isolate different apoptotic blebs [39,40]. According to their size, apoptotic blebs can be classified as apoptotic bodies (ApoBD, 1-5 µm) or apoptotic microvesicles (MVs, 0.1-1 µm) [41]. Using a series of centrifugation steps, we separated ApoBD and MVs from the final cleared supernatant (SN) of dexamethasone-treated thymus homogenates (Fig 5C). Next, we analyzed the presence of Ccl6, Ccl9/10, Ccl12, Ccl21 and Cxcl10 in the three fractions by Western blot. As shown in Fig 5D, regardless of their PS-binding activity, each chemokine was found to some extent in the ApoBD and MV fractions as well as in the final SN fraction. However, chemokines and cytokines might be transported on the surface or inside the vesicles [42]. Since suitable mouse chemokine antibodies were not available to probe vesicles by FACS, we turned to functional assays, reasoning that since liposomal PS binding did not interfere with chemokine bioactivity (Fig 3) then, apoptotic vesicle surface-bound but not encapsulated chemokines should activate cognate GPCRs. As expected, an appropriate control agonist and SN fraction triggered calcium flux responses in L1.2 cell lines expressing Ccr1 (receptor for the non-PS-binding Ccl6, Ccl9/10), Ccr2 (the receptor for Ccl12) and Cxcr3 (the receptor for Cxcl10). In contrast, when purified ApoBD and MV fractions, each containing the same chemokines as SN, were used as stimuli, calcium flux responses were induced only in Ccr2- and Cxcr3-expressing L1.2 cells, not in Ccr1-expressing cells (Fig 5E). Importantly, when the ApoBD fraction was filtered to remove apoptotic bodies, the Ccr2-mediated calcium response was nearly eliminated (Panel A in S9 Fig). Filtration of the SN fraction, however, did not affect activation of Ccr2-expressing cells (Panel A in S9 Fig). Similarly, preincubation of the ApoBD fraction with a neutralizing anti-Ccl12 antibody, but not with an irrelevant control antibody, blunted the Ccr2-mediated calcium flux signal (Panel B in S9 Fig). These results indicated that the apoptotic body-Ccl12 complex is required for the activation by the ApoBD fraction of Ccr2, a chemokine receptor typically expressed in phagocytes. To assess Ccl21 activity in the fractions, since our Ccr7 reporter cell line responded poorly in calcium flux assays but well in chemotaxis assays to recombinant Ccl21, we turned to this latter type of assay. As expected, SN induced chemotaxis of both Ccr1- and Ccr7-expressing cells (Fig 5F). However, only Ccr7-transfected cells migrated towards the ApoBD and MV fractions (Fig 5F). The reduced chemotactic activity observed with the ApoBD fraction compared to the MV fraction could be explained by the larger size of ApoBD, which may impede diffusion through the transwell filter and limit access to the reporter cells. These results indicate that active PS-binding chemokines are transported on the surface of apoptotic blebs.

**Fig 5.**
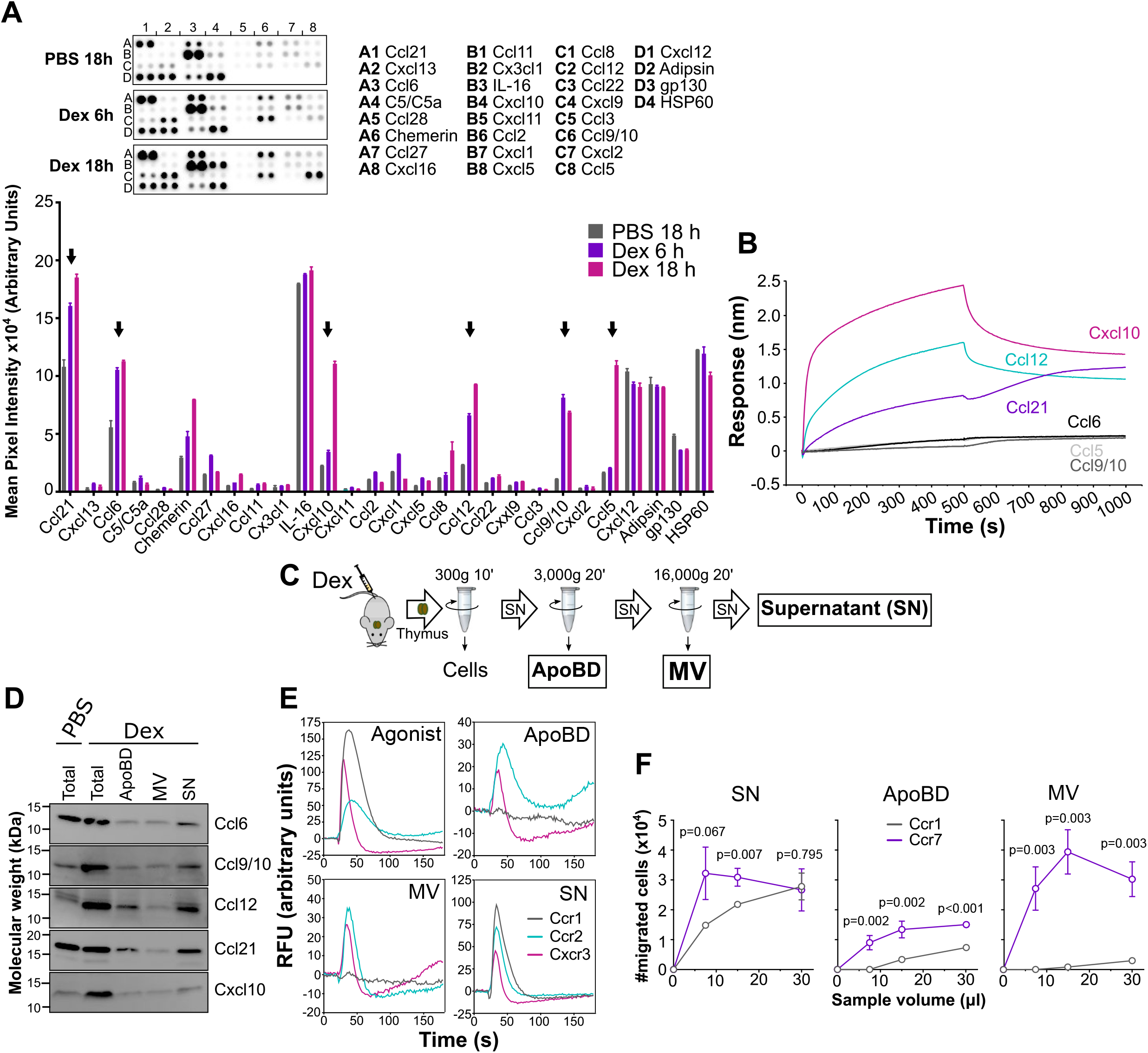
Endogenous phosphatidylserine-binding chemokines bound to apoptotic blebs in vivo activate cognate GPCRs. A) Endogenous chemokine expression in the thymus of dexamethasone (Dex)-treated mice. Top, anti-chemokine antibody array membranes were incubated with 200 µg of total protein of mouse thymus extracts collected 6 or 18 h after i.p. injection with PBS or Dex. The membranes are spotted in duplicate with capture antibodies for the chemokines indicated on the right side (position coordinates are indicated with lettered rows and numbered columns). Positions D2 (Adipsin), D3 (gp130) and D4 (HSP60) correspond to loading controls. Bottom, mean ± SD of the mean pixel intensity for each chemokine. Black arrows point to highly expressed chemokines. B) BLI analysis of the binding of mouse Ccl5, Ccl6, Ccl9/10, Ccl12, Cxcl10 and Ccl21 (500 nM) to DOPS liposomes. Final binding sensorgrams were generated after subtraction of the binding recorded on DOPC sensors used as reference. C) Schematic representation of the centrifugation steps followed to isolate apoptotic bodies (ApoBD), microvesicles (MV) and the final cleared supernatant (SN) from thymus homogenates of Dex-treated mice. D) Western blot analysis of Ccl6, Ccl9/10, Ccl12, Ccl21 and Cxcl10 in ApoBD, MV and SN fractions isolated from thymus 18 h after i.p. injection of mice with PBS or dexamethasone. “Total” lane corresponds to the supernatant obtained right after pelleting the cells. E and F) Functional assays using ApoBD, MV and SN fractions isolated from mouse thymus 18 h after i.p. injection of dexamethasone. E) Calcium flux assays using L1.2 reporter cell lines expressing Ccr1, Ccr2 and Cxcr3 (as indicated in the inset of the “SN” panel). Panel “Agonist” shows the calcium flux response after addition of 50 nM of a known ligand for each receptor (Ccl9/10 for Ccr1, Ccl12 for Ccr2, and Cxcl10 for Cxcr3). Calcium recordings correspond to the mean of duplicates from one experiment representative of 3 independent experiments. F) Chemotaxis assays using reporter cell lines expressing Ccr1 and Ccr7 (as indicated in the inset of the “ApoBD” panel) and increasing volumes (x axis) of the indicated fractions. Data are the mean ± SD of the number of migrated cells of triplicates from one experiment representative of 3 independent experiments. p values from multiple t tests corrected for multiple comparisons by the Holm-Sidak method are indicated for each volume data point.

## Discussion

In this study, we identify a previously unappreciated and broadly shared capacity of chemokines to bind anionic phospholipids and provide proof of principle for PS-bound chemokine action as find-me signals during the elimination of apoptotic cells. PS binding was chemokine-specific and high affinity, and was observed when PS was presented either in pure form, or in the context of liposomes and biological membranes from apoptotic cell lines, primary cells and extracellular vesicles derived from apoptotic primary cells. Importantly, PS-bound chemokines retained their ability to activate cognate GPCRs on leukocytes. Although our study began with a chemokine phospholipid screen, we have focused our attention on the interaction of chemokines with PS. Future work will be needed to analyze the biochemistry and biological significance of chemokine binding to other anionic phospholipids.

Since most chemokines are highly basic proteins, it is not surprising that they bind to highly anionic molecules like PS and GAGs. More surprising is the degree of specificity revealed by our biochemical screen, which suggested that chemokine-PS interaction is not exclusively charge-driven. First, we showed that basic chemokines such as CXCL8 (pI = 9.1), do not bind PS. Second, we found that most chemokines displayed a null/weak binding to phosphoinositides, which are more highly anionic phospholipids than PS. Furthermore, as demonstrated by BLI, many chemokine-PS interactions displayed a very slow dissociation rate, suggesting a contribution of hydrophobic or other stable bonds in the formation of the chemokine-PS complex. Importantly, not all GAG-binding chemokines were able to interact with PS. In particular, we showed that mouse and human CCL5, which are strong GAG-binders [32,43], do not bind PS. Furthermore, while soluble heparin neutralized CCL20 and CCL21, these chemokines remained fully active in the presence of very high doses of DOPS liposomes. Additionally, we showed that the elimination of CCL20-DOPS liposomes by pull-down reduced the overall chemotactic activity. These results suggest that the chemokine GAG- and PS-binding sites may require different molecular determinants and may map differently relative to the receptor-binding site. It is important to note that like GAGs, different chemokines may bind PS in different ways, and therefore, case-by-case and comprehensive analyses may be required to fully understand the structural basis of chemokine-PS interaction.

In preliminary experiments, we found that thymocytes undergoing apoptosis in vivo switch chemokine tethering mechanisms, downregulating anionic GAGs from live cells and upregulating anionic PS on apoptotic cells, in both cases promoting presentation of chemokines to leukocyte GPCRs. Additional work will be needed to assess how general this phenomenon may be. GAGs act as substrates for the formation of haptotactic gradients promoting binding and oligomerization of chemokines on cell surfaces [5]. In fact, receptor-binding chemokine mutants unable to oligomerize or interact with GAGs are inactive in vivo [32]. Like GAGs, PS is mostly found on cell surfaces, and here we demonstrate that it can also induce chemokine oligomerization. Interestingly, we show that PS exposure is accompanied by a massive depletion of GAGs on the surface of apoptotic thymocytes, which bind chemokines in an AnV- and MFG-E8-susceptible manner. These two PS-binding proteins, however, did not block chemokine binding to live GAG-competent thymocytes, which lack PS exposure on the cell surface. We found that most of the chemokines tested, with the exception of CXCL11, displayed a stronger binding to live thymocytes than to apoptotic thymocytes. Therefore, we propose that whereas in the presence of GAGs chemokines might preferentially bind to GAGs over PS, PS may substitute for GAGs as a mediator of chemokine oligomerization and haptotactic gradients in tissues with high apoptotic rates. It is important to note that tumor cells and virally-infected cells also expose PS [44–47], therefore, chemokine modulation by PS may become more apparent in non-homeostatic conditions.

Effective and rapid apoptotic cell clearance is essential for the proper maintenance of homeostasis. Phagocytes use chemotactic find-me signals and the surface PS eat-me signal to reach and recognize apoptotic cells, respectively [48]. Surprisingly, to date, chemokines, arguably the most potent chemoattractants of the immune system, are only represented by CX3CL1 in the list of reported find-me signals, which instead include highly atypical chemotactic cues such as apoptotic blebs. Several independent studies have reported that apoptotic bodies and microvesicles released by apoptotic cells can induce chemotaxis of phagocytes [25–27]. However, how these extracellular vesicles exert their chemotactic action had remained unknown. Here we show that while many chemokines can be found in apoptotic bodies and microvesicles, these apoptotic blebs only activate cognate GPCRs of PS-binding chemokines. These results support a simple explanation: Apoptotic blebs are chemotactic because they are coated with chemokines tethered to surface-exposed PS, and therefore, they can act as find-me signals thanks to their complex with PS-binding chemokines. Consistent with this, we demonstrate that binding to PS-containing liposomes does not affect the chemokine activity. We propose a mechanism by which, to reach the dying cell, phagocytes would follow a trail of extracellular vesicles presenting chemokines on their surface (Fig 6). Apoptotic cells are known to produce macrophage-attracting chemokines [49], but chemokines could also be released by neighboring healthy cells. However, live cell-derived chemokines might be expressed at lower levels and might not attract phagocytes. In addition, GAGs might retain chemokines on the surface of live cells (Fig 6). In contrast, we show here that cells lose surface GAGs during apoptosis, and therefore, in the absence of GAGs, chemokines released by apoptotic cells would readily interact with PS on the surface of apoptotic cells and apoptotic blebs to establish a gradient of chemokine-presenting microvesicles able to guide phagocytes toward the dying cell (Fig 6). In addition, vesicle-bound phagocyte-recruiting and inflammatory chemokines could be rapidly eliminated through phagocytosis of the vesicle by phagocytes (Fig 6), contributing to the overall anti-inflammatory nature of apoptosis and collaborating with atypical chemokine receptors in the regulation of chemokine availability in the extracellular compartment [50]. This mechanism would potentially extend the classification of find-me signal to any PS-binding chemokine expressed by apoptotic cells able to activate GPCRs expressed in phagocytes like CCR2. Our study provides proof of principle of this notion exemplified in the case of Ccl12. Here we show that Ccl12 expression is induced in mouse apoptotic thymus, that it binds PS and that the complex of thymus apoptotic blebs with mouse Ccl12 is required to activate mouse Ccr2-expressing cells. In addition, we found that other CCR2 ligands in humans, CCL2 and CCL13, also interact with PS. Although, we did not observe a significant induction of Ccl2 in mouse apoptotic thymus, the chemokine expression profile of apoptotic cells may differ among tissues or cell types. For instance, it has been previously shown that Ccl2 upregulated in mouse lung by bleomycin-induced injury enhances the removal of apoptotic neutrophils [51]. Furthermore, it is likely that different chemokines are expressed in other modalities of cell death with more highly inflammatory conditions (necrosis, necroptosis…), which would result in the attraction of different subsets of phagocytes. A comprehensive analysis of the chemokines and chemokine receptors expressed by dying cells and phagocytes, respectively, would help to identify other chemokine-apoptotic body-receptor axes involved in the recruitment of phagocytes and to understand their influence on the kinetics and efficiency of dying cell clearance.

**Fig 6.**
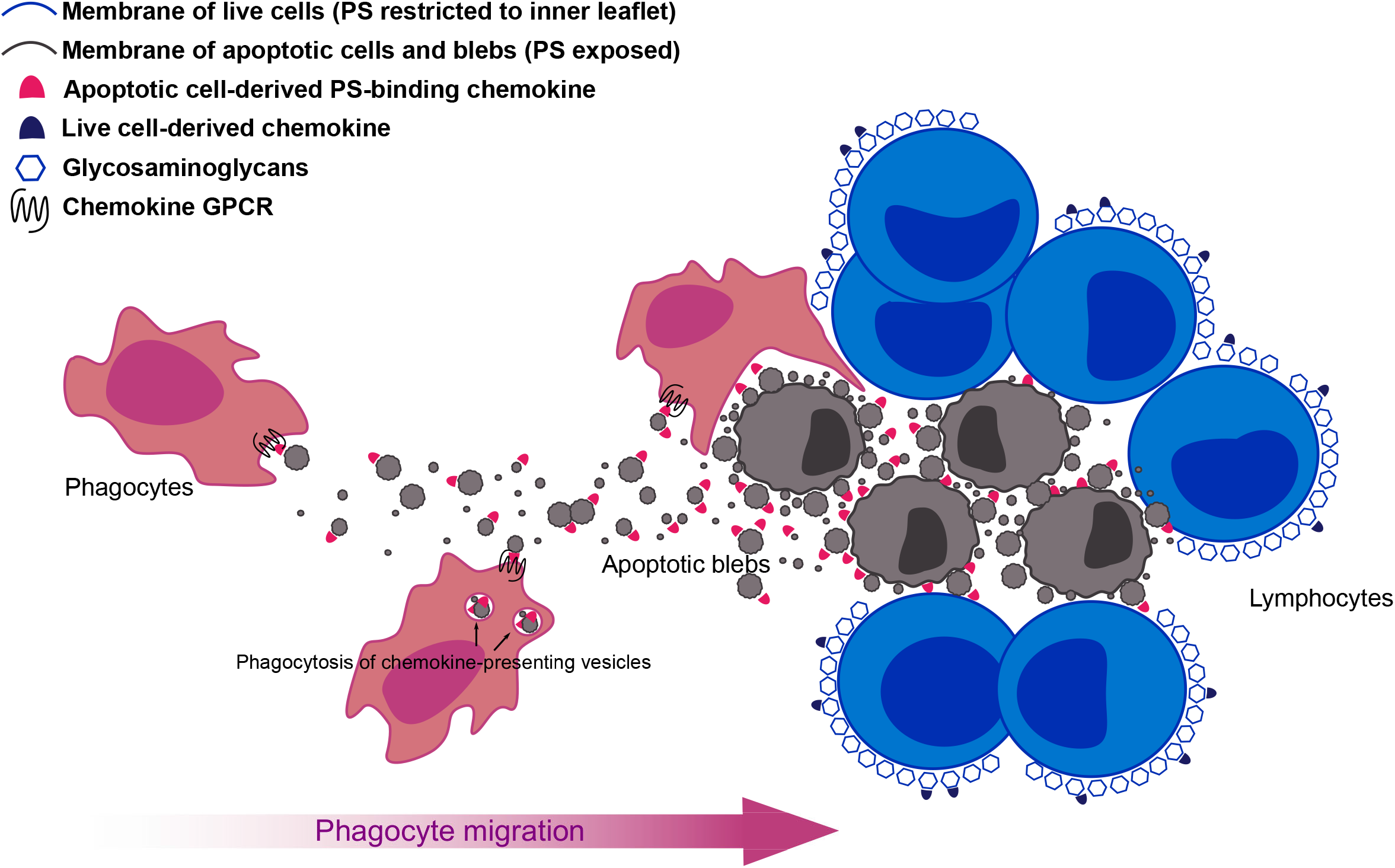
A gradient of chemokine-presenting apoptotic blebs guide phagocytes toward apoptotic cells for their elimination. Graphic representation of the proposed model for the role of PS-binding chemokines in apoptotic cell clearance. Chemokines produced by live cells are retained by GAGs on the cell surface whereas in the absence of GAGs in apoptotic cells, phagocyte-recruiting chemokines interact with PS exposed on the surface of apoptotic cells and blebs to form a gradient for phagocyte recruitment. Phagocytes engulf and eliminate chemokine-presenting apoptotic blebs and apoptotic cells reducing inflammation.

Here we chose to validate the biological relevance of chemokine-PS interactions in the context of apoptosis. However, many PS-binding chemokines identified by our study are not known to recruit phagocytes and therefore, they are unlikely to impact apoptotic cell clearance. Instead, PS-binding may regulate the activity of chemokines that attract non-phagocytic cell types in other biological processes where cell membrane PS becomes exposed and accessible for the interaction with soluble cytokines, like cancer and infectious diseases. Extracellular vesicles like apoptotic bodies, microvesicles and exosomes have received increasing attention in recent years as essential elements of intercellular communication in cancer and immunity [52,53]. RNA transcripts, microRNA, enzymes, cytokines, chemokines and cellular receptors, among others, are transported from cell to cell by extracellular vesicles [54]. Research in the field has focused principally on encapsulated nucleic acids and integral membrane proteins of the vesicle. This might be a consequence of the fact that while it is easy to understand how cell state can be modulated by RNAs, cellular receptors or cytosolic enzymes delivered by extracellular vesicles, how vesicle-encapsulated cytokines and chemokines, whose action depend on the interaction with receptors in the plasma membrane of a target cell, might contribute to vesicle-mediated cell-to-cell communication is less apparent. In this regard, our study provides proof of principle that signaling cytokines can be displayed and distributed in vivo attached to the surface of extracellular vesicles. Tumor cells and viruses might use this system to distribute cytokines and activate or antagonize cellular receptors in distant organs. Understanding how lipids interact with secreted proteins may help explain the functional roles played by extracellular vesicles during homeostasis, cancer, and infection.

## Materials and methods

### Animals

C57BL/6j mice were obtained from The Jackson Laboratory (Bar Harbor, ME). All mice were maintained under specific pathogen–free housing conditions at an American Association for the Accreditation of Laboratory Animal Care–accredited animal facility at the National Institute of Allergy and Infectious Diseases (NIAID) and housed in accordance with the procedures outlined in the Guide for the Care and Use of Laboratory Animals under the protocol LMI-8E approved on 31/12/2015 and annually renewed by the Animal Care and Use Committee of NIAID.

### Cells

CHO-K1 and the GAG-deficient variant CHO-745 cells (generous gift from Dr. Antonio Alcami, Spanish Research Council, Spain) were grown in DMEM/F12-Glutamax (Life Technologies, Carlsbad, CA) supplemented with 10% FBS. L1.2 cells (kindly provided by Dr. Eugene Butcher, Stanford University, CA) were grown in RPMI-Glutamax (Life Technologies) supplemented with 10% FBS, 1 mM sodium pyruvate and 0.1 mM non-essential amino acids. CHO-K1, CHO-745 and L1.2 cells were cultured at 37°C and 5% CO_2_. Suspension Expi293F cells were maintained in FBS-free Expi293 Expression Medium (both from Life Technologies) and grown at 37°C, 8% CO_2_ with constant agitation at 125 rpm.

### Reagents

Untagged human/mouse recombinant chemokines produced in *E. coli* were obtained from Peprotech (Rocky Hill, NJ). Of note, recombinant CXCL16 included only the extracellular domain of human CXCL16. The synthetic biotinylated human chemokines CCL2, CCL3, CCL11, CXCL11 and CCL20 were purchased from Almac (Craigavon, UK). All primary anti-chemokine antibodies used in this study were purified polyclonal rabbit antibodies from Peprotech. Recombinant human MFG-E8 protein produced in mouse myeloma cells was purchased from R&D Systems (Minneapolis, MN).

### Generation of stable cell lines expressing chemokine receptors

Cell lines expressing the mouse chemokine receptors Ccr1, Ccr2, Ccr6, Ccr7 and Cxcr3 were generated in L1.2 cells. cDNA clones in pCMV6 plasmid for these chemokine receptors were obtained from Origene (Rockville, MD). 2 million L1.2 cells were transfected with 2 µg of plasmid using the SG Cell Line transfection kit and a 4D-Nucleofector X (both from Lonza, Walkersville, MD) following the manufacturer’s instructions. 48 h post-transfection, cells were seeded in 24-well plates in pools containing 2,400 cells/well in selection media (RPMI-Glutamax, 10% FBS, 1 mM sodium pyruvate, 0.1 mM non-essential amino acids and 1 mg/ml geneticin). After 7 days at 37°C, media was replaced with fresh selection media and the cells were incubated for 7 additional days. Then, surviving pools were screened for the expression of the corresponding chemokine receptor by calcium-flux assays. Pools displaying the strongest calcium flux responses were further selected by limiting dilution in 96-well plates. Receptor-expressing clones were selected by calcium flux assays. Highly responsive clones were expanded and frozen in geneticin-free selection media containing 5% DMSO. For subsequent experiments, cell lines were grown in selection media.

### Lipid array binding assays

The lipid binding specificity of recombinant human chemokines (Peprotech) was determined using Membrane Lipid Strips obtained from Echelon BioSciences (Salt Lake City, UT). These strips are spotted with 100 pmol of 15 different lipids found in cell membranes. First, strips were incubated with blocking buffer (1X Tris-buffered saline [TBS], 0.1% Tween-20, 3% BSA) for 1h at room temperature. Then, recombinant chemokines were added at 0.1 µg/ml in blocking buffer and incubated for 1 h at room temperature. Strips were washed with TBS-T (TBS, 0.1% Tween-20) and bound protein was detected with specific rabbit anti-chemokine antibodies (Peprotech) followed by an HRP-conjugated anti-rabbit antibody (Abcam, Cambridge, MA). Strips were developed with WesternBright Sirius (Advansta, San Jose, CA) and imaged in an Omega Lum C imager (Gel Company, San Francisco, CA).

### Liposomes

The phospholipids 1,2-dioleoyl-sn-glycero-3-phosphocholine (DOPC), 1,2-dioleoyl-sn-glycero-3-phospho-L-serine (DOPS), 1,2-dioleoyl-sn-glycero-3-phosphoethanolamine-N-(cap biotinyl) (DOPEbiot), and 1,2-distearoyl-sn-glycero-3-phosphoethanolamine-N-[biotinyl(polyethylene glycol)-2000] (DSPE-PEGbiot) were all purchased from Avanti Polar Lipids (Alabaster, AL). In this study, two different types of liposomes were used: “DOPC”, composed of DOPC:DOPEbiot at a 95:5 ratio (weight %), and “DOPS”, composed of DOPS:DOPC:DOPEbiot at a 30:65:5 (weight %). Phospholipids resuspended in chloroform were mixed at the indicated ratios (1 mg total lipid) and the solvent was evaporated in a SpeedVac Concentrator (Fisher Scientific, Pittsburgh, PA). Dried lipids were rehydrated in 0.5 ml of PBS or TBS for 2 h at room temperature. Subsequently, large unilamellar vesicles were prepared by extrusion (> 11 passes) through a 0.1 µm membrane at room temperature using a mini-extruder (Avanti Polar Lipids). Liposomes were used immediately after preparation.

### Liposome binding assays

The ability of chemokines to interact with phospholipid liposomes was tested by ELISA and BLI. For ELISA-based binding experiments, DOPC and DOPS liposomes prepared in TBS were immobilized at 10 µg/ml in binding buffer (TBS, 0.1% BSA) on streptavidin-coated high capacity plates (Thermo Fisher, Waltham, MA). Increasing concentrations (0.01-10 µg/ml) of recombinant chemokines in binding buffer were incubated in the wells for 15 min at room temperature. Plates were washed extensively with TBS and bound chemokine was detected with specific rabbit anti-chemokine antibodies (Peprotech) and HRP-conjugated anti-rabbit secondary antibody development (Abcam). The (Twin)-Strep-tag /Strep-Tactin affinity purification system (IBA LifeSciences, Göttingen, Germany) was used to produce twin-strep-tagged fusion chemokines which were detected with an HRP-conjugated anti-Streptag mAb (IBA LifeSciences). Plates were developed with TMB One Component (Surmodics, Eden Prairie, MN) and the reaction was stopped with sulfuric acid before measuring the absorbance at 450 nm (A_450_) in a FlexStation 3 microplate reader (Molecular Devices, Sunnyvale, CA). Non-specific binding to the plate was corrected by the subtraction of the A_450_ recorded for each chemokine concentration in wells incubated without liposome. Where indicated, chemokine binding to liposome-coated wells was competed by preincubation of chemokines with increasing concentrations of lipids for 30 min at room temperature.

BLI experiments were performed in an Octet RED384 system (Pall ForteBio, Fremont, CA) essentially as previously described with some modifications [55]. For these experiments, liposomes were prepared in PBS and DSPE-PEGbiot instead of DOPEbiot was used as biotinylated phospholipid to improve the immobilization of the liposomes on the biosensors. Briefly, streptavidin-coupled biosensors pre-wetted in PBS for at least 10 min were equilibrated in PBS for 60 s. Then, DOPC and DOPS liposomes were immobilized to a final 2-4 nm or 0.5-1 nm response for binding screening or kinetic purposes, respectively. Subsequently, biosensor tips were washed for 60 s in PBS and blocked for 300 s with 0.05% BSA in PBS. Finally, biosensors were washed for 150 s in PBS and the association of recombinant chemokines was recorded for 500 s followed by a 500 s dissociation in PBS. All the steps were performed at 30° C and 1,000 rpm. Final binding sensorgrams were generated by subtracting the binding to biosensors loaded with DOPC liposomes from the binding recorded in DOPS biosensors. For binding screening, chemokines were tested at 1 µM. For kinetic analysis, increasing concentrations (50 nM to 1,000 nM) of chemokines were used and sensorgrams were adjusted to a 1:1 Langmuir model.

### Chemokine-phospholipid cross-linking assays

The capacity of different natural phospholipids — brain phosphatidylethanolamine (PE), brain phosphatidylcholine (PC), brain phosphatidylserine (PS) and heart cardiolipin (CL), all from Avanti Polar lipids — to induce chemokine oligomerization was studied by crosslinking experiments. First, phospholipids stored in chloroform at 10 mg/ml were diluted 1:1000 in 20 mM HEPES. Then, 50 ng of chemokine were incubated for 30 min at room temperature in the presence or absence of phospholipid at a 1:8 (chemokine:total lipid) molar ratio in 12 µl of 20 mM HEPES. Then, 3 µl of the crosslinker BS_3_ (Life Technologies) were added for a 0.25 mM final concentration and samples were incubated at room temperature for additional 30 min. The reaction was stopped by the addition of 5 µl of 4X SDS-PAGE loading buffer and chemokine oligomerization was analyzed by Western blot. Chemokine oligomers were detected by specific rabbit anti-chemokine antibodies (Peprotech) followed by development with an HRP-conjugated anti-rabbit antibody (Abcam).

### Chemotaxis assays

The effect of DOPC and DOPS on the activity of human CCL3, CCL20 and CCL21 was assessed by transwell chemotaxis assays. For this, the indicated chemokine concentrations were preincubated with increasing molar ratios (10^2^-10^5^) of total lipid of DOPC or DOPS liposomes for 30 min at room temperature in chemotaxis buffer (DMEM-Glutamax [Life Technologies] supplemented with 0.1% of BSA and 10 mM HEPES). To confirm chemokine binding to the liposomes under these conditions, 50 µl of the 1:10^4^ chemokine:liposome mix was incubated with streptavidin-coated plates and the liposome-bound chemokine was analyzed by ELISA as explained above. Also, where indicated, chemokines were preincubated with a 10^3^-fold molar excess of soluble heparin (Sigma, St. Louis, MO). Chemokine samples were added to the bottom chambers of 96-well ChemoTx plates (Neuroprobe, Gaithersburg, MD) and 200,000 L1.2 cells expressing appropriate receptors (Ccr1 for CCL3, Ccr6 for CCL20 and Ccr7 for CCL21) were added in chemotaxis buffer to the top chambers and allowed to migrate through a 5 µm pore-sized membrane for 3-4 h at 37° C. Then, the top chamber and filter were removed, 5 μl/well of CellTiter 96 Aqueous One Solution (Promega, Madison, WI) was added to the bottom wells and the absorbance at 490 nm (A_490_) was determined after incubation for 2 h at 37° C using a FlexStation 3 plate reader (Molecular Devices). The total number of cells that migrated to the bottom wells was calculated by interpolation of the observed A_490_ values using a standard curve generated with a known number of cells.

### Chemokine binding to apoptotic cells

Apoptotic CHO-745 cells and mouse thymocytes were used for chemokine binding experiments. Apoptosis of GAG-deficient CHO-745 cells was induced by irradiation with UV light. Cells cultured on 100 mm plates were washed once with PBS and cell monolayers were irradiated with 100 mJ in a UV-Stratalinker 2400 (Stratagene, San Diego, CA) with the plate lid off or on to generate the mock control. 6 h after treatment, cells were collected with trypsin-EDTA followed by at least two washes with culture media to remove the EDTA, which may interfere with the Ca^2+^-dependent binding of AnV. Apoptotic thymocytes were generated by incubation of thymocytes (10^7^ cells/ml) freshly isolated from C57BL/6 mice with 1 µM dexamethasone for 4-5 h in RPMI-glutamax supplemented with 10% FBS. All binding assays were performed in AnV Binding Buffer (Biolegend, San Diego, CA). For competition experiments, cells were preincubated for 5 min at room temperature with 1 µg of the unlabeled PS-binding proteins MFG-E8 (R&D systems) and AnV (Biolegend). 300,000 cells were incubated with 250 nM biotinylated CCL3, CCL11, CXCL11 (all from Almac, Craigavon, UK), or AnV (Biolegend) in 100 μl of binding buffer for 10 min at room temperature. Then, cells were pelleted by centrifugation (1,200 rpm, 5min) and incubated in 100 μl of binding buffer containing 0.125 μg of streptavindin-APC (Biolegend) for 10 min at room temperature. Finally, cells were resuspended in binding buffer and stained with 3.5 µg/ml propidium iodide (Biolegend), and, in the case of the thymocytes, also with 0.1 µM YO-PRO (Life Technologies). 30,000 events were acquired in an LSR Fortessa cell analyzer and FlowJo software (both from Becton Dickinson, San Jose, CA) was used for the data analysis.

### Chemokine expression in mouse thymus

The expression of chemokines in the thymus of C57BL/6j mice injected (i.p.) with PBS or 250 µg of dexamethasone (Sigma) was analyzed using Proteome Profiler Mouse Array kit (R&D systems). 6 h or 18 h after treatment, mouse thymus was collected and homogenized in 1 ml of cold PBS containing cOmplete protease inhibitor cocktail (Roche, Indianapolis, IN) using a tissue homogenizer (Omni International, Kennesaw, GA). Protein concentration was determined by BCA assay and array membranes were incubated with an extract volume corresponding to 200 µg of protein. Arrays were developed following the manufacturer’s instructions and membranes for all samples were imaged simultaneously in an Omega LumC imager (Gel Company). The mean pixel intensity for each signal was analyzed using ImageJ.

### Functional assays using apoptotic bleb fractions from mouse thymus

The chemokine activity retained in mouse apoptotic blebs was analyzed by calcium flux and chemotaxis assays. C57BL/6j mice were injected i.p. with PBS or 250 µg of dexamethasone. In each experiment, 2 mice were treated with PBS and 3 with dexamethasone. 18 h after treatment, thymi were collected and pooled together for each treatment. Thymi were homogenized using 70 µm cell strainers and 500 µl/thymus of assay buffer. For calcium flux assays thymus samples were prepared and tested in calcium assay buffer (Cell culture media:Calcium buffer [10% FBS RPMI-Glutamax:HBSS supplemented with 20 mM HEPES] mixed 1:1 vol:vol), and for chemotaxis assays thymi were homogenized in chemotaxis buffer (DMEM-Glutamax supplemented with 0.1% BSA and 10 mM HEPES). Then, samples were processed by a series of centrifugation steps as shown in Fig 5C. First, intact cells were removed by centrifugation for 10 min at 300xg. The supernatant was collected and recentrifuged at 3,000xg for 20 min. The resulting pellet was resuspended using the same volume of assay buffer and defined as ApoBD (apoptotic bodies), while the supernatant was recentrifuged at 16,000xg for 20 min. The supernatant of this last step was collected and defined as SN (supernatant) and the pellet was resuspended in the same volume of assay buffer and defined as MV (microvesicles). All centrifugation steps were performed at 4° C. Fractions were then used in calcium flux or chemotaxis assays using L1.2 cells stably transfected with Ccr1, Ccr2, Cxcr3 or Ccr7. For calcium flux assays, 300,000 cells/well in black clear-bottom plates were incubated with Calcium 6 reagent (Molecular Devices) for 2 h at 37° C. Calcium flux signals were recorded for 180 s in a FlexStation 3 plate reader (Molecular Devices). For this, 25 µl/well of each fraction or the same volume containing 50 nM of the appropriate recombinant agonist diluted in calcium assay buffer were dispensed using the reader’s pipettor robot at 20 s after the initiation of the recording. For chemotaxis assays, different volumes of each fraction were tested essentially as explained above.

## Acknowledgments

This work was supported by the Division of Intramural Research of the National Institute of Allergy and Infectious Diseases, National Institutes of Health.

## Author Contributions

Conceptualization, S.M.P and P.M.M; Investigation and analysis, S. M. P; Writing-original draft, S.M.P; Writing-review and editing, S.M.P and P.M.M; All authors have read and agreed to the published version of the manuscript.

## Supporting Information

See attached files.

