## Supplementary Figures for "Chemokines as phosphatidylserine-bound ‘find-me’ signals in apoptotic cell clearance"

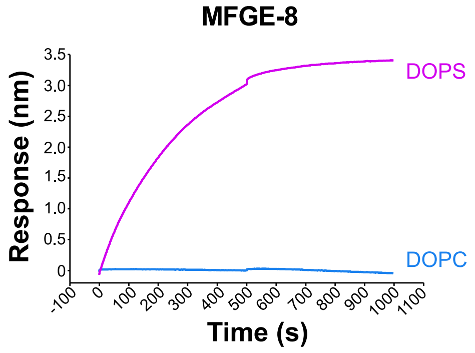


**S1 Fig. Biolayer interferometry assays can be used to analyze the phosphatidylserine-binding activity of recombinant proteins.**

Binding of recombinant MFG-E8 (200 nM) to BLI biosensors immobilized with pure DOPC (blue sensorgram) or DOPS (pink sensorgram) liposomes.

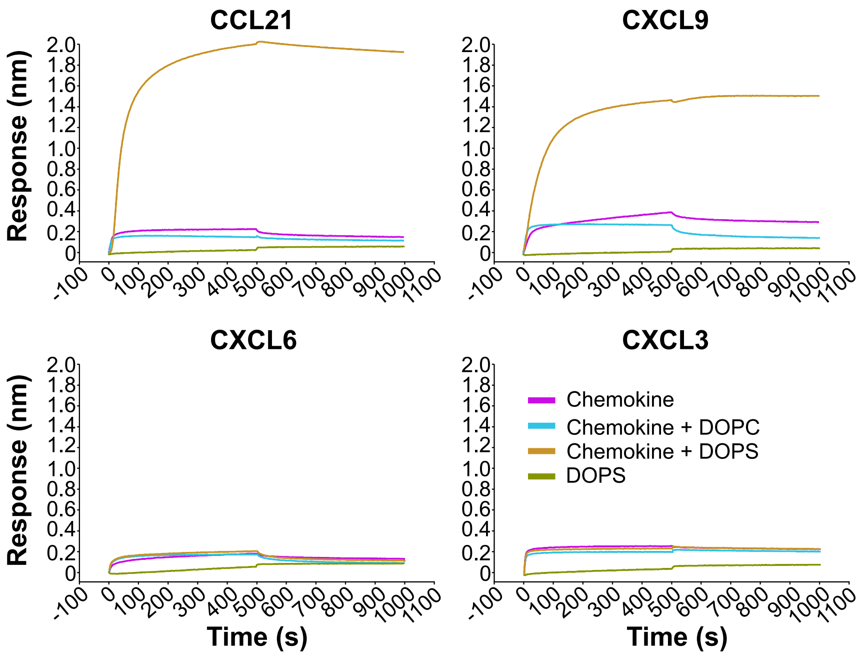


**S2 Fig. Some primary anti-chemokine antibodies may not recognize their target chemokine when in complex with DOPS liposomes.**

Primary antibodies used in ELISA or protein-lipid overlay assays to detect CCL21, CXCL9, CXCL6 or CXCL3 were immobilized onto BLI amine-reactive biosensors. BLI binding sensorgrams for the interaction of DOPS liposomes alone (green) and 400 nM of each chemokine alone (magenta) or preincubated with 0.5 mg/ml of DOPS (yellow) or DOPC (blue) liposomes are shown. Increase in the binding response in the presence of liposomes is indicative of the binding of a large analyte (chemokine-liposome complex).

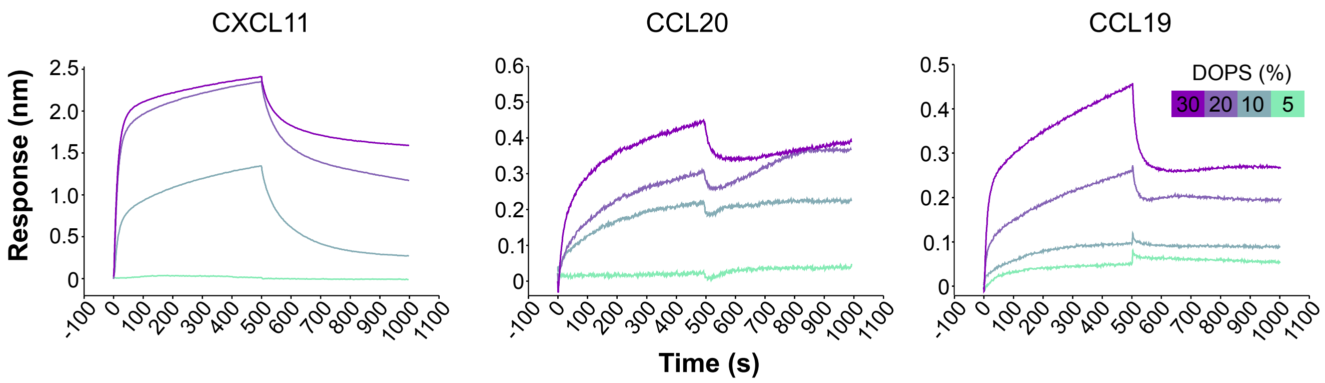


**S3 Fig. Chemokines detect 5-10% of PS in liposomes.**
BLI experiments showing the binding of the indicated chemokines (500 nM) to DOPC liposomes containing decreasing amounts of DOPS (as indicated in the inset of the CCL19 graph). Binding to pure DOPC liposomes was subtracted from all binding curves.

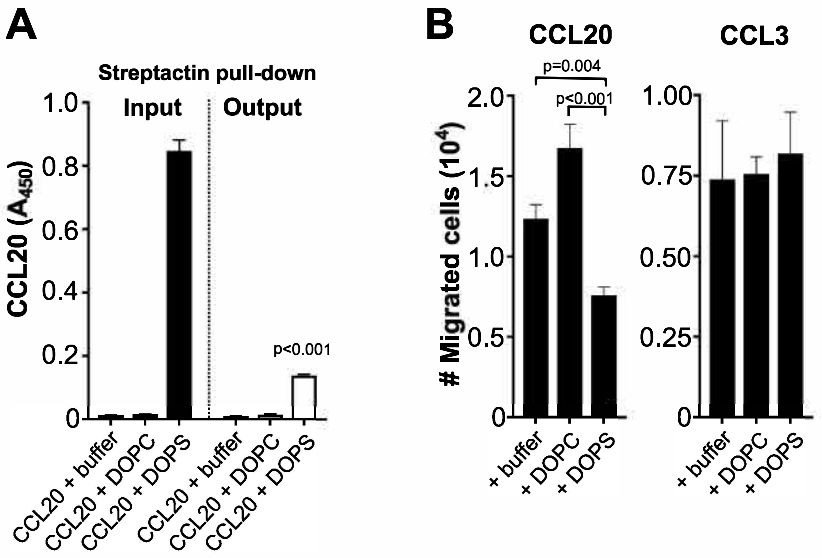


**S4 Fig. Depletion of CCL20-DOPS liposome complexes by pull-down reduces cell migration.**

A) Pull-down of CCL20-DOPS liposome complexes decreases their availability in solution. CCL20 (1 nM) was incubated with buffer or a 104-fold molar excess of DOPC or DOPS liposomes. 50 µl before (input) and after (output) pull-down with 30 µl of Strep-Tactin beads were analyzed in triplicates by ELISA in streptavidin-coated plates. Liposome-bound CCL20 was detected with a rabbit anti-CCL20 pAb followed by an HRP-conjugated anti-rabbit antibody and absorbance at 450 nm (A450) was determined after developing with TMB One Component solution. Bars represent mean SD from one experiment representative of two independent experiments. The p value from a two-tailed t test for the analysis of CCL20 + DOPS input vs output is indicated. B) Pull-down of CCL20-DOPS liposome complexes reduces cell migration. CCL3 or CCL20 (as indicated above each graph) were incubated with buffer or a 104-fold molar excess of DOPC or DOPS liposomes. Then, chemokine-liposome complexes were pull down with Strep-Tactin beads and the chemokine activity in the supernatants was tested by chemotaxis assays using L1.2 cell lines expressing the appropriate chemokine receptor (Ccr6 for CCL20 and Ccr1 for CCL3). Bars represent mean SD of the number of migrated cells of triplicates from one experiment representative of 3 independent experiments. p values are from one way ANOVA with Bonferroni correction for multiple comparisons.

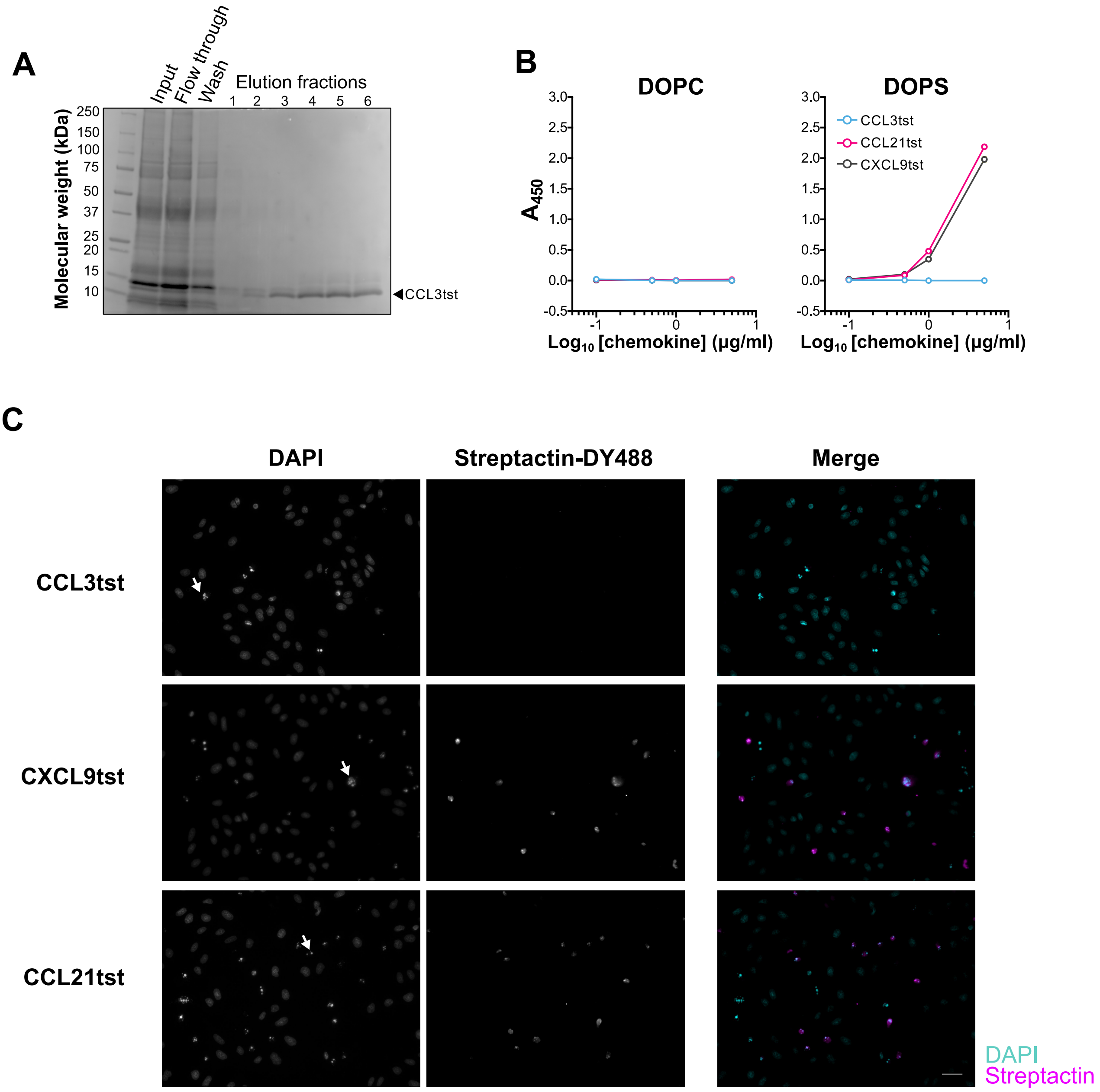


**S5 Fig. Glycosylated recombinant chemokines interact with phosphatidylserine and apoptotic cells.**

A) Purification of recombinant CCL3 expressed in Expi293F cells. Coomasie-stained acrylamide gel showing the purification steps of human CCL3. Human CCL3, CXCL9, and CCL21 were tagged with a twin-strep tag (tst) at the C-terminus and expressed in Expi293F cells. Recombinant proteins were purified from cell supernatants by affinity chromatography using Strep-Tactin XT columns. B) CXCL9tst and CCL21tst interact with PS-containing liposomes. The binding of in-house produced chemokines to DOPC or DOPS liposomes was analyzed by ELISA. Increasing doses (x axis) of the different chemokines (as indicated in the inset of the right panel) were incubated in wells immobilized with DOPC (left panel) or DOPS (right panel) liposomes. Wells were washed profusely with TBS, and bound chemokine was detected with an HRP-conjugated anti-tst mAb and absorbance at 450 nm (A450) was determined after developing with TMB One Solution substrate. Data are the mean ± SD of triplicates from one experiment representative of 3 independent experiments. C) PS-binding chemokines interact with the surface of dying CHO-745 cells. Immunofluorescence images showing the binding of CCL3tst, CXCL9tst and CCL21tst (as indicated on the left side of each row) to UV-irradiated CHO-745 cells. Cells cultured on coverslips were exposed to 100 mJ of UV light using a Stratalinker. 6h after treatment, coverslips were incubated with 400 nM of each chemokine in AnV binding buffer. After washing, samples were stained with Strep-Tactin XT conjugated with DY-488, fixed and mounted using Prolong Gold with DAPI. Epifluorescence images were acquired at 20x in a Zeiss Anxiovert 200M inverted microscope. DAPI (first column), Strep-Tactin staining (second column) and merge images (third column) are shown. White arrows in the DAPI panels point at examples of dying cells displaying fragmented and condensed nuclei. In the merge panels, DAPI and Strep-Tactin staining are shown in cyan and magenta, respectively, as indicated on the bottom right corner. A white scale bar corresponding to 50 µm is shown in the merge CCL21tst panel.

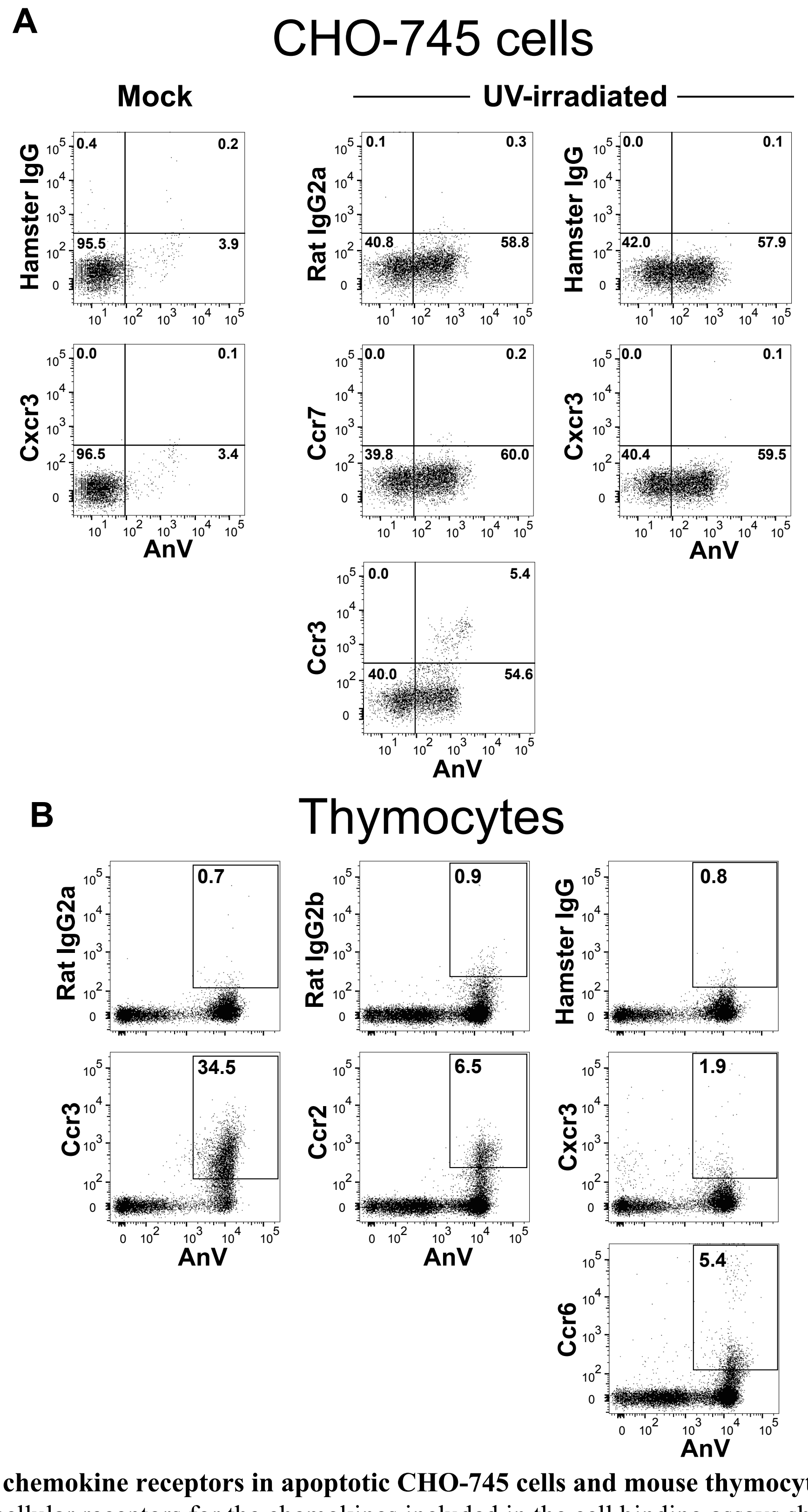


**S6 Fig. Expression of chemokine receptors in apoptotic CHO-745 cells and mouse thymocytes.**

The expression of the cellular receptors for the chemokines included in the cell binding assays shown in Figure 4 was analyzed by FACS in CHO-745 cells (A) and mouse thymocytes (B) as indicated above each panel. A) Cxcr3, Ccr3 and Ccr7 are not expressed in live or apoptotic CHO-745 cells. Mock-treated or UV-irradiated CHO-745 cells (as indicated above the graph columns) were stained with annexin V (AnV)-APC and PE conjugated antibodies for the chemokine receptors Ccr3, Ccr7 and Cxcr3 (as indicated on the y-axis of the corresponding graphs). Chemokine receptor-AnV dot plots are shown. Dot-plots for the staining with the pertinent isotype controls are shown above the corresponding columns. B) Ccr3, and to a lower extent, Ccr2 and Ccr6, but not Cxcr3, are expressed specifically by apoptotic thymocytes. Freshly isolated mouse thymocytes were incubated with 1 μM dexamethasone at 37 C. After 4 h, thymocytes were stained with AnV-APC and PE conjugated anti-Ccr2, anti-Ccr3, anti-Ccr6 or anti-Cxcr3 antibodies as indicated on the y-axis of the corresponding graphs. Chemokine receptor-AnV dot plots are shown below the corresponding isotype control for each anti-chemokine receptor antibody. In both, panel A and B, numbers indicate % of events in each gate.

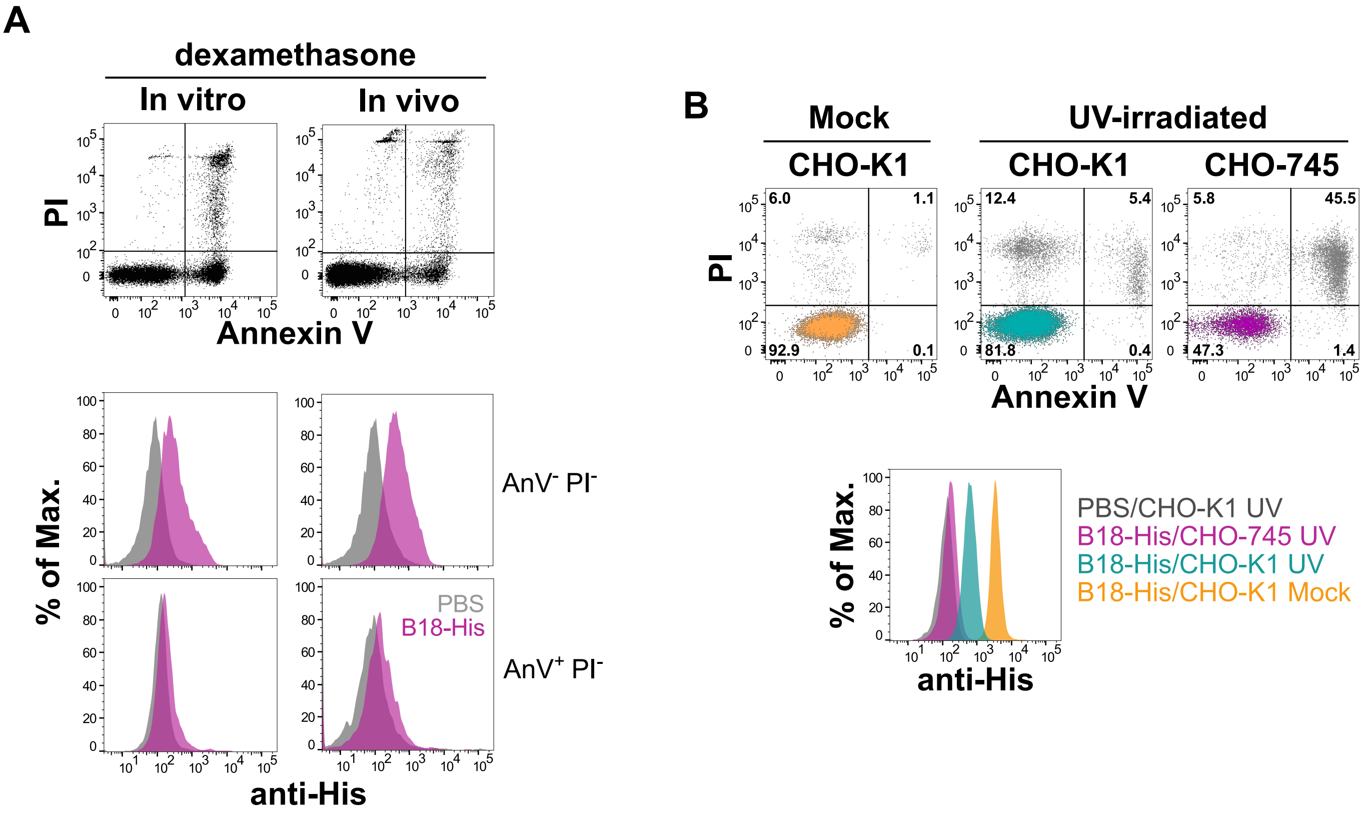


**S7 Fig. Apoptotic cells display highly reduced levels of surface GAGs.**

A) GAGs are depleted from the surface of apoptotic thymocytes. Top graphs show dot-plots for the Annexin V (AnV) and PI staining of mouse thymocytes treated with dexamethasone in vitro or in vivo, as indicated above each graph column. Mouse apoptotic thymocytes were generated in vitro by incubating freshly isolated thymocytes with 1 µM dexamethasone at 37 C, or in vivo by injecting 250 µg of dexamethasone in C57BL/6j mice via i.p. Thymocytes were collected 6 h after treatment and incubated with PBS alone (gray histograms) or 200 nM of a His-tagged B18 protein (B18-His, purple histograms) as indicated in the inset of the bottom right graph. Bound protein was detected with an anti-His mAb followed by an anti-mouse Alexa Fluor 488-conjugated antibody. Before the analysis, cells were stained with PI and AnV-APC. Bottom graphs represent histograms for the binding of B18 to the surface of live (AnV- PI-) and early apoptotic cells (Annexin V+ PI-), as indicated on the right side of each graph row. B) CHO-K1 cells exposed to UV-light present diminished levels of surface GAGs. CHO-K1 (GAG competent) and CHO-745 (GAG deficient) cells were exposed to 100 mJ of UV-light. 6 h after irradiation, cells were collected and the binding of B18-His was analyzed as in A. Top dot plots show the PI and AnV staining of mock- and UV-irradiated CHO-K1 or CHO-745 cells as indicated above each graph. Numbers correspond to % of events in each gate. Bottom histogram graph shows the binding of B18-His to AnV- PI- populations (color coded in the top dot plots) from mock- (orange) or UV-irradiated CHO-K1 (green) and CHO-745 cells (purple) as indicated in the legend on the right side of the graph. Staining of UV-irradiated CHO-K1 cells in the absence of B18-His (PBS, gray) is shown as reference.

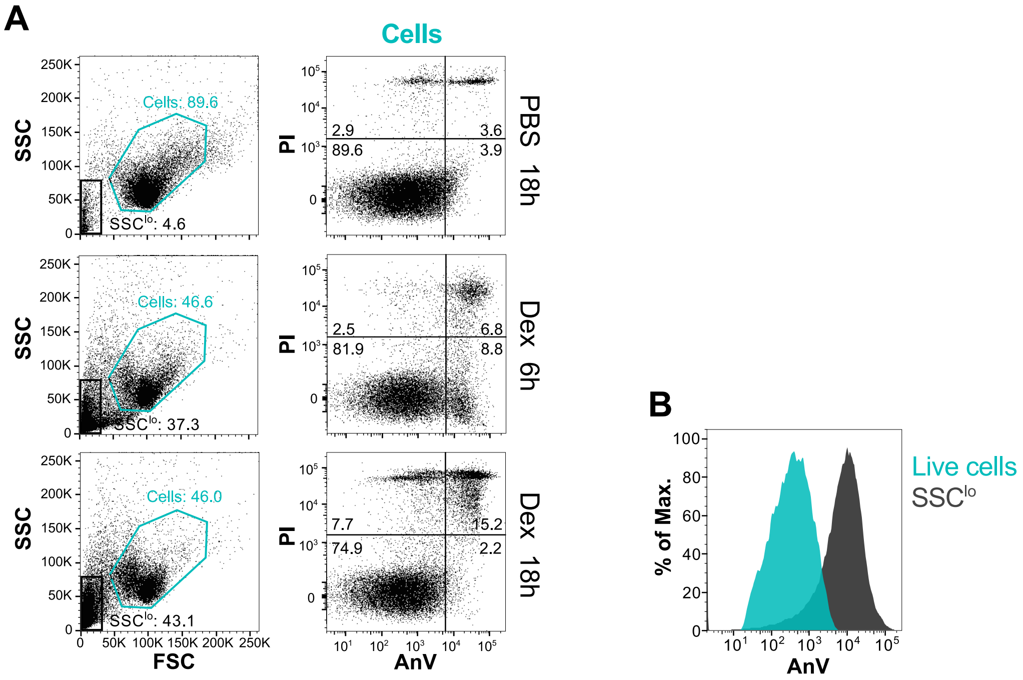


**S8 Fig. Injection of dexamethasone induces the production of PS-containing apoptotic blebs in mouse thymus.** A) C57BL/6j mice were i.p. injected with PBS or dexamethasone (Dex) and thymocytes were isolated 6 or 18 h after treatment (as indicated on the right side of each graph row) and stained with propidium iodide (PI) and annexin V (AnV)-APC. In the left column, SSC-FSC dot plots showing gates for the cells (Cells, blue) and apoptotic blebs (SSClo, black). In the right column, PI-AnV dot plots of the events from the "Cells" gate of each condition. Numbers indicate % of events in each gate. B) Histograms are for the AnV staining of live cells (AnV-PI-) and apoptotic blebs (SSClo, black).

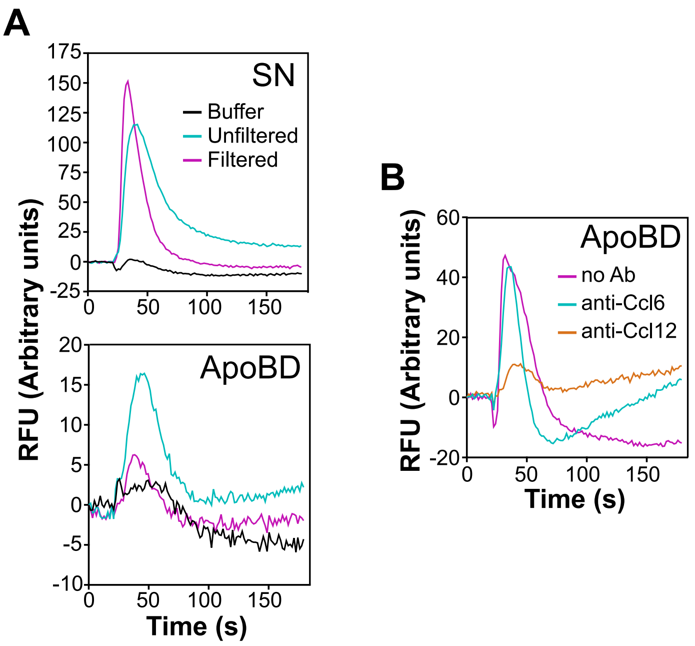


**S9 Fig. Apoptotic body-Ccl12 complex is required for Ccr2 activation by ApoBD fraction.**

A and B) Calcium flux assays using Ccr2-expressing L1.2 cells, and ApoBD and SN fractions isolated from mouse thymus 18 h after i.p. injection of dexamethasone. A) Calcium flux response for buffer alone (black) and for the mouse SN and ApoBD fractions previously filtered (pink) or not (blue) through a 0.2 m filter. B) Calcium flux response obtained with the ApoBD fraction preincubated or not (No Ab, pink) with 10 g/ml of an anti-Ccl12 antibody (orange) or a control anti-Ccl6 antibody (blue). Calcium recordings in A and B correspond to the mean of duplicates from one experiment representative of 2 independent experiments.
