## Supplementary Tables for "Chemokines as phosphatidylserine-bound ‘find-me’ signals in apoptotic cell clearance"

**S1 Table.** Binding response (second column) of human chemokines (first column) at 1 µM to DOPS liposomes by biolayer interferometry. Binding fold change relative to the binding of the known PS-binding chemokine CXCL16 is indicated in the third column. Binding responses < 0.1 nm are highlighted in orange. Fold change values > 2.0 and < 0.5 are highlighted in blue or purple, respectively.

| | Binding at 495 s (nm) | Fold change |
|---|---|---|
| **CXCL16** | 0.428 | 1.000 |
| **CCL2** | 0.407 | 0.951 |
| **CCL3** | 0.041 | 0.096 |
| **CCL11** | 1.346 | 3.145 |
| **CCL13** | 1.554 | 3.631 |
| **CCL17** | 0.278 | 0.650 |
| **CCL19** | 1.158 | 2.706 |
| **CCL20** | 0.660 | 1.542 |
| **CCL21** | 0.312 | 0.729 |
| **CCL22** | 0.247 | 0.577 |
| **CCL23** | 0.061 | 0.143 |
| **CCL24** | 0.146 | 0.341 |
| **CXCL1** | 0.194 | 0.453 |
| **CXCL2** | 1.559 | 3.643 |
| **CXCL3** | 0.348 | 0.813 |
| **CXCL4** | 2.396 | 5.598 |
| **CXCL5** | 0.140 | 0.327 |
| **CXCL6** | 1.867 | 4.362 |
| **CXCL8** | 0.055 | 0.129 |
| **CXCL9** | 1.142 | 2.668 |
| **CXCL11** | 2.912 | 6.804 |
