## Supplementary Methods for "Chemokines as phosphatidylserine-bound ‘find-me’ signals in apoptotic cell clearance"

### Supporting materials and methods

##### Expression and purification of recombinant chemokines

Synthetic codon-optimized genes encoding CCL21, CXCL9 and CCL3 were obtained from Life Technologies. A twin-strep tag (TST, seq: SAWSHPQFEKGGGSGGGSGGSAWSHPQFEK) was added at the immediate C-terminus of each chemokine for purification and detection purposes (IBA Lifesciences, Göttingen, Germany). Genes were subcloned into a pcDNA3.1 plasmid for expression of the proteins under a CMV promotor. For the expression of recombinant protein, 150 $x10^6$ Expi293F cells were transfected with 60 μg of plasmid using the Expifectamine 293 Transfection Kit (Life Technologies) following the manufacturer's instructions. 4-7 days after transfection, protein secreted into the supernatant was purified by affinity chromatography using Strep-TactinXT gravity columns (IBA Lifesciences) following the manufacturer's guidelines. Briefly, cell-free supernatant was buffered with 1X Buffer W (100 mM Tris pH 8.0, 150 mM NaCl, 1 mM EDTA) and biotin in the culture media was blocked with 18.1 ml/L of BioLock Solution (IBA LifeSciences). Subsequently, the supernatant was further clarified by centrifugation (4,000 rpm, 15 min) and applied at a 1 ml/min rate into a Strep-TactinXT gravity column. The column was washed with 5 ml Buffer W and bound protein was eluted with 2 ml Buffer BXT (100 mM Tris pH 8.0, 150 mM NaCl, 1 mM EDTA, 50 mM biotin). Protein-containing elution fractions were pooled together, and concentrated and dialyzed with PBS using 3 kDa Amicon Ultra 0.5 ml Centrifugal Filters (Millipore, Bedford, MA). Protein concentration was determined by SDS-PAGE and densitometry, and proteins were stored at -80° C.

##### Pull-down of chemokine-liposome complexes

To investigate the relative contribution of free chemokines versus chemokines complexed with liposomes to cell migration, we tested chemotactic activity remaining in solution after depletion of chemokine-liposome complexes by specific pull-down. For this, 1 nM CCL3 or CCL20 was incubated with buffer or a $10^4$-fold molar excess of DOPC or DOPS liposomes in 0.4 ml of chemotaxis buffer (DMEM-Glutamax supplemented with 0.1% BSA and 10 mM HEPES) for 30 min at room temperature. Then, a total of 30 µl of Strep-TactinXT-coupled agarose beads (IBA LifeSciences), previously washed 5 times with chemotaxis buffer, was added to each sample and incubated with rotation for 30 min at room temperature. Beads were pelleted by centrifugation (12,000 rpm, 1 min) and the supernatants were collected and tested by ELISA and chemotaxis assays as explained previously.

**Analysis of the expression of chemokine receptors in apoptotic cells**

The expression of chemokine receptors of interest in apoptotic CHO-745 cells and mouse thymocytes was analyzed by FACS. Apoptosis was induced in each cell type as described previously. 300,000 cells were first incubated with TruStain FcX in PBS-staining buffer (1X PBS, 1% BSA and 1% FBS) to block Fc receptors. Then, cells were stained with one of the following PE conjugated antibodies (all from Biolegend) in PBS-staining buffer for 20 min on ice: anti-Ccr2 (clone SA203G11), anti-Ccr3 (clone J073E5), anti-Ccr6 (clone 29-2L17), anti-Ccr7 (clone 4B12), anti-Cxcr3 (clone CXCR3-173), rat IgG2a isotype control, rat IgG2b isotype control or armenian hamster IgG isotype control. Subsequently, cells were washed with Annexin V (AnV) binding buffer (Biolegend) and stained with AnV-APC (Biolegend). 30,000 cells were acquired in a LSRFortessa cell analyzer and analyzed using FlowJo (both from Becton Dickinson).

**Analysis of surface GAGs in apoptotic cells**

The levels of GAGs on the surface of apoptotic mouse thymocytes and CHO cells were determined by evaluating the cell binding activity of B18 by FACS. B18 is a soluble interferon binding protein encoded by vaccinia virus that interacts with very high affinity (nanomolar range) with cell surface GAGs (including heparan sulfate and chondroitin sulfate). Apoptosis of mouse thymocytes and CHO-K1 (GAG-competent) or CHO745 (GAG-deficient) cells was induced by dexamethasone treatment or UV light irradiation, respectively, as explained previously. 300,000 cells were incubated with PBS or 200 nM recombinant his-tagged B18 protein (kind gift from Dr. Alcami, Spanish Research Council) in PBS-staining buffer (1X PBS, 1% BSA and 1% FBS) for 20 min on ice. Then, cells were washed and stained with a mouse anti-PentaHis antibody (Qiagen, Germantown, MD) followed by an AlexaFluor 488-conjugated anti-mouse antibody. Subsequently, cells were washed with AnV binding buffer and stained with AnV-APC and propidium iodide (both from Biolegend). 30,000 cells were acquired in a LSRFortessa cell analyzer and analyzed using FlowJo (both from BD).
